# The competition between splicing and 3′ processing shapes the human transcriptome

**DOI:** 10.1101/2025.08.19.671063

**Authors:** Lindsey V. Soles, Shuangyu Li, Liang Liu, Kristianna S.K. Sarkan, Erik G. Alvstad, Lusong Tian, Yoseop Yoon, Marielle Cárdenas Valdez, Ivan Marazzi, Yongsheng Shi

## Abstract

Eukaryotic pre-mRNA processing steps, including splicing and 3′ processing, are tightly coordinated, yet the underlying mechanisms remain incompletely understood. U1 snRNP has been proposed to inhibit 3′ processing at intronic polyadenylation (IPA) sites through a splicing-independent mechanism termed telescripting. In contrast, we discovered that disrupting splicing—by targeting various key components such as U1 snRNP, U2 snRNP, U2AF, and SF3b—activates 3′ processing at thousands of IPA sites. Notably, splicing inhibition, especially of U1 snRNP, induced widespread premature transcription termination within gene bodies through both IPA-coupled and -independent mechanisms. Different splicing factors activated overlapping and distinct sets of IPA sites, reflecting their specific contributions to transcription and spliceosome function. Conversely, inhibition of 3′ processing enhanced splicing globally. These findings support a model in which splicing and 3′ processing are competing processes that intersect with transcription to shape the transcriptome landscape.

## INTRODUCTION

The maturation of RNA Polymerase II (RNAPII) transcripts involves several processing steps, including 5′ capping, splicing, and 3′ processing. Splicing takes place in the spliceosome, which is composed of five snRNP complexes and numerous proteins.^1,2^ The 5′ splice site (ss) and the branch point are recognized by base-pairing with the U1 and U2 snRNAs, respectively, while the polypyrimidine tract and 3′ ss are bound by the U2AF2 and U2AF1 subunits of the U2AF complex.^3^ The spliceosome is assembled in a stepwise manner, transitioning from E, to A, then B, and ultimately to C complex in which the splicing reactions occur.^2,3^ Pre-mRNA 3′ processing typically entails an endonucleolytic cleavage followed by the addition of a poly(A) tail in a process called cleavage and polyadenylation (CPA).^4,5^ Most mammalian poly(A) sites (PASs) include an A(A/U)UAAA hexamer, a downstream U/GU-rich element, and additional auxiliary sequences.^4,5^ These sequences are recognized by the CPA machinery, including CPSF and CstF.^6^ Once the CPA complex is assembled, the pre-mRNA is cleaved by the endonuclease CPSF73 and the poly(A) tail is added by poly(A) polymerase.^7,8^ In recent years, pre-mRNA processing has emerged as a potential therapeutic target for many diseases, including cancer. A number of small molecule inhibitors have been identified for both splicing and 3′ processing. For example, Spliceostatin A (SSA) and Pladienolide B (PB) inhibit splicing by binding to SF3b, a subcomplex of the U2 snRNP.^9,10^ Another compound, OTS964, blocks splicing by inhibiting the kinase CDK11,^11^ which is responsible for phosphorylating SF3b1.^12^ JTE-607 and a class of benzoxaboroles block 3′ processing by inhibiting the endonuclease CPSF73.^13–16^ Some of these inhibitors display antitumor activities and have been proposed as potential candidates for cancer therapy.^11,16–20^

Many splicing and 3′ processing events occur co-transcriptionally^21,22^ and these two processes are tightly coordinated to ensure accurate and efficient gene expression. Such coordination could be positive or negative. For example, splicing of an upstream intron has long been known to promote 3′ processing and this activity is mediated by direct interactions between U2 snRNP and the CPSF complex.^23,24^ Additionally, the poly(A) tail has been shown to stimulate splicing, especially of the last intron.^25,26^ On the other hand, although approximately 30% of annotated PASs are found in introns,^27^ most intronic PASs are repressed. Repression of intronic polyadenylation (IPA) is important because IPA transcripts often encode truncated and non-functional proteins.^28,29^ Aberrant IPA site usage has been proposed to inactivate tumor suppressor genes in leukemia^28^ and DNA repair genes in prostate and ovarian tumors.^30^ Previous studies provided evidence that IPA sites are repressed by U1 snRNP bound at the 5′ ss or within introns via a splicing-independent mechanism.^31,32^ This model, called telescripting, was based on the observation that blocking base-pairing interactions between U1 snRNA and pre-mRNA with an antisense morpholino oligo (AMO) broadly activates IPA site usage.^31,32^ In contrast, a U2 snRNA-targeting AMO or a small molecule splicing inhibitor SSA displayed less of an effect on IPA site usage.^31^ However, later global analyses demonstrated that blocking splicing with small molecule inhibitors, such as PB, or targeting U4 snRNA with an AMO did activate IPA to varying degrees.^33–35^ Further, treatment with a U1 snRNA-targeting AMO has recently been shown to induce premature transcription termination, which could indirectly modulate IPA site usage.^36^ To reconcile these conflicting models, it is important to systematically characterize the relationship between splicing and 3′ processing.

In this study, we address the question of how splicing and 3′ processing impact each other. We inhibited splicing using six different methods and found that all treatments broadly activated IPA. Different splicing-inhibiting treatments activated both overlapping and distinct IPA sites and such specificity was determined, at least in part, by alterations in RNAPII elongation and termination caused by these treatments. Conversely, we found that inhibiting 3′ processing by using JTE-607 improved splicing efficiency globally. In contrast to the previously proposed splicing-independent mechanism, our results support a model in which splicing and 3′ processing compete co-transcriptionally and the outcome of this competition determines the transcriptome. These findings are clinically relevant as splicing inhibitors activate IPA and consequently lead to decreased expression of the full-length transcripts of many genes, including tumor suppressor genes, potentially representing a serious risk in using general splicing inhibitors for cancer therapy.

## RESULTS

### Global splicing inhibition activates intronic polyadenylation

To systematically investigate the relationship between pre-mRNA splicing and 3′ processing, we disrupted splicing in HEK293T cells using six different methods and evaluated their impact on pre-mRNA 3′ processing (Fig. 1A). As described earlier, the 5′ ss, the branch point, and the 3′ ss are recognized by U1 snRNP, U2 snRNP, and U2AF respectively to assemble the A complex, which subsequently transitions to form B complex, then B^act^, and finally the C complex in which the splicing reactions take place.^3^ To inhibit splice site recognition, we treated cells with antisense morpholino oligos (AMOs) against U1 or U2 snRNAs, which block their base-pairing with the 5′ ss and branch point respectively, or depleted U2AF1 and U2AF2 by RNAi (Fig. 1A and S1A). Additionally, we treated cells with the small molecule PB, which inhibits the U2 snRNP component SF3b1 and prevents the transition from A to B complex.^10,37,38^ Lastly, we treated cells with a CDK11 inhibitor, OTS964, that blocks splicing at a later step by preventing the transition from B to B^act^ complex.^12^ Following these treatments, RNA was extracted from these cells and subjected to PAS-seq analyses. PAS-seq is a 3′-end RNA sequencing technique that precisely maps polyadenylation site usage and measures polyadenylated RNA levels genome-wide (Fig. S1B).^39,40^

**Figure 1.**
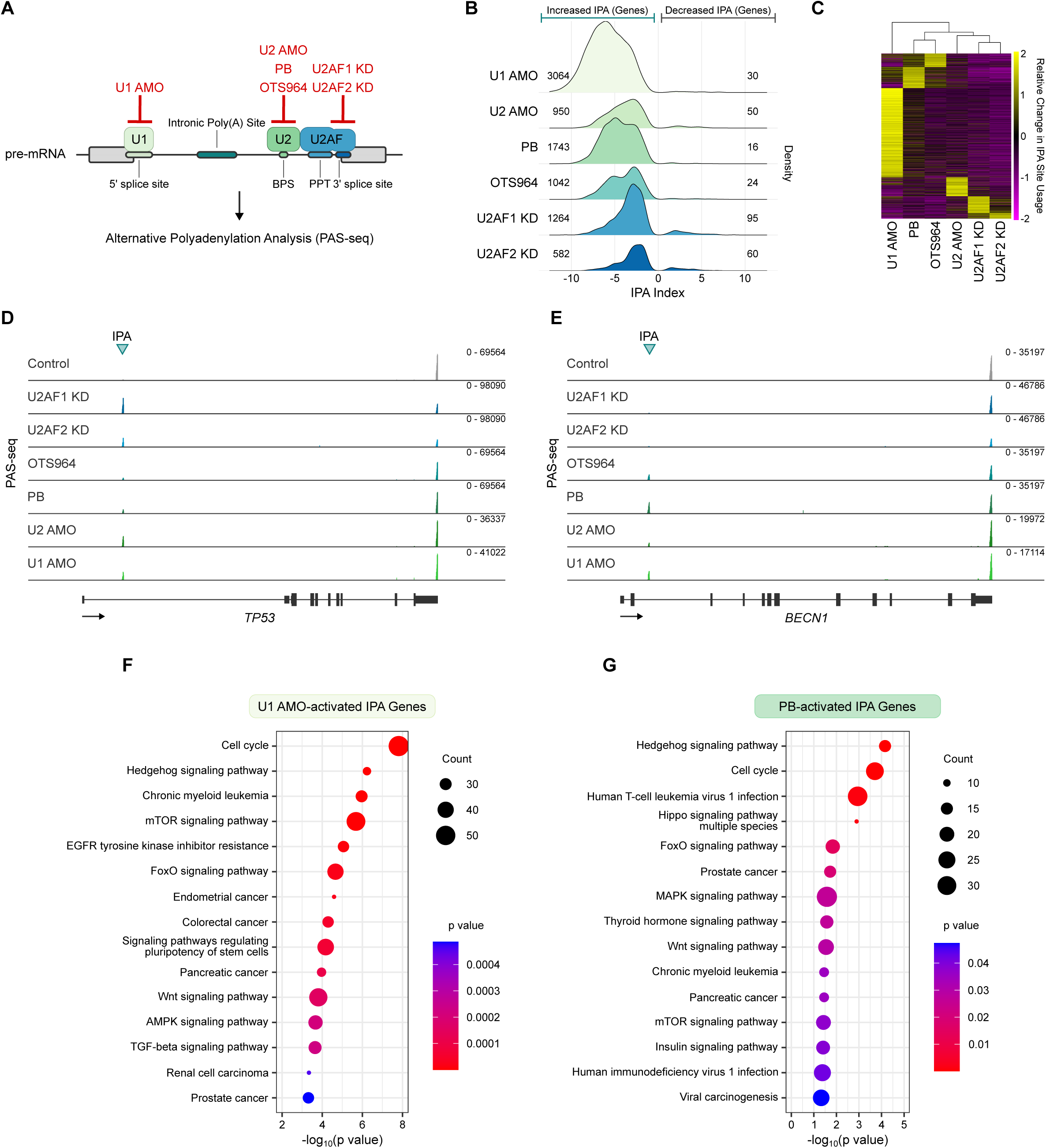
Global splicing inhibition activates intronic polyadenylation. **(A)** Schematic of the global splicing inhibitions tested followed by PAS-seq. **(B)** Ridgeline density plots depicting the change in IPA site usage following all global splicing inhibitions tested. Values to the left of 0 indicate genes with increased IPA site usage. Values to the right of 0 indicate genes with decreased IPA site usage.

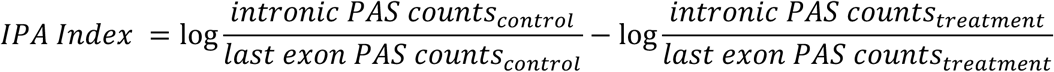 **(C)** Heatmap of the relative change in IPA site usage scaled by row across all treatments. A positive value (yellow) indicates that the IPA site is activated upon splicing inhibition. Hierarchical clustering was performed across treatments. **(D-E)** PAS-seq data tracks for the genes *TP53* (**D**) and *BECN1* (**E**) following the indicated splicing inhibition treatments. Each peak represents reads mapped to the 3′ end of mRNAs. **(F-G)** KEGG pathway analyses for activated IPA sites following treatment with U1 AMO(**F**) or PB (**G**). The terms related to cell development, growth and disease were collected. The terms in which p value < 0.05 and FDR < 0.05 were regarded as a significant functional annotation clustering.

Consistent with previous reports,^31,32^ we observed robust and widespread activation of IPA following U1 AMO treatment with 3,064 genes showing significantly higher IPA usage (FDR < 0.05 and increase by more than 15% of total transcripts) while only 30 genes displayed reduced IPA (FDR < 0.05 and decrease by more than 15% of total transcripts) (Fig. 1B). Interestingly, however, we found that all other splicing inhibition treatments also led to widespread increases in IPA and very few changes in the opposite direction (Fig. 1B and Fig. S1B-C). These results are in contrast with previous reports, which suggested that U1 AMO, but not other splicing inhibitors, specifically activates IPA in a splicing-independent manner.^31^ Instead, our data provided strong evidence that global splicing inhibition by individually targeting distinct splicing factors leads to widespread activation of IPA.

We next compared the IPA sites that were activated by different splicing inhibiting treatments. A heatmap for all IPA sites that displayed significantly increased usage in at least one condition demonstrated that some IPA sites were activated by all splicing inhibitors (Fig. 1C). This trend was evident in two example genes, *NECAP2* and *CCDC71* (Fig. S1D-E). In both cases, very low levels of IPA transcripts were detected in control cells. However, all six treatments led to significantly increased levels of IPA transcripts and concomitant decreases in the full-length transcripts, suggesting that splicing inhibition activated IPA in these genes (*NECAP2*: 20.6-73.2% increase in relative IPA site usage, *CCDC71*: 34.1-87.3% increase in relative IPA site usage). Hierarchical clustering of these IPA changes revealed the closest clustering for treatments that targeted the same splicing factor or complex (Fig. 1C). For example, OTS964 and PB, both of which inhibit SF3b1 functions, clustered together (Fig. 1C). Similarly, depletion of U2AF1 and U2AF2 clustered together (Fig. 1C). This U2AF cluster was also closely related to U2 snRNP inhibition by U2 AMO (Fig. 1C). Importantly, splicing inhibiting treatments also activated IPA in tumor suppressor genes including *TP53* and *BECN1* (Fig. 1D-E) (*TP53*: 22.1-64% increase in relative IPA site usage, *BECN1*: 15.7-26.8% increase in relative IPA site usage).^41–43^ Indeed, ontology analyses of genes with activated IPA following splicing inhibition revealed significant enrichment within the cell cycle, cancers, and cancer-related signaling pathways (Fig. 1F-G and Fig. S1F-H). Taken together, these results strongly suggest that global splicing inhibition by individually targeting multiple distinct splicing factors led to broad activation of IPA, including within tumor suppressor genes, and that inhibiting different splicing factors activated overlapping and distinct IPA sites.

### Gene-specific splicing inhibition activates intronic polyadenylation

Our observation that global splicing inhibition induced IPA activation can be explained by a competition between splicing and 3′ processing. Alternatively, it has been previously suggested that blocking splicing globally could lead to sequestration of U1 snRNP on unspliced pre-mRNAs and such a decrease in the concentration of free U1 snRNP could in turn result in activation of IPA due to diminished inhibition by U1 snRNP via the telescripting mechanism.^34^ To distinguish between these possibilities, we inhibited splicing of individual introns and determined its impact on IPA. If IPA is repressed in a splicing-dependent manner, we would predict that intron specific-disruptions of 5′ or 3′ ss recognition would activate IPA. On the other hand, if IPA is repressed in a U1 snRNP-dependent and splicing-independent manner, blocking the 3′ ss of an individual intron should have no effect on IPA. To test this, we generated stable cell lines that expressed a series of minigene reporters in a doxycycline-inducible manner and monitored the spliced or IPA transcript levels by 3′ Rapid Amplification of cDNA Ends (3′ RACE) (Fig. 2A and Fig. S2A). The initial reporter included an IPA-containing intron and its flanking exons (exons 13 and 14) from the *KPNB1* gene (Fig. 2A and Fig. S2B). This IPA site in the *KPNB1* gene was activated by U1 AMO treatment (Fig. S2B). Under control conditions, only fully spliced RNAs, but no IPA transcripts, were detected for the *KPNB1* reporter (Fig. 2B, lane 2), similar to the transcripts of the endogenous gene (Fig. S2B). However, U1 AMO treatment led to robust IPA site usage (49.2%, Fig. 2B, lane 3), again displaying a similar pattern to the transcripts of the endogenous gene (Fig. S2B). We next mutated the 5′ or 3′ ss to weaker non-consensus sequences and tested the impact of lower splicing efficiency on IPA site usage (Fig. 2A and Fig. S2C). When the 5′ ss was mutated, nearly half of the reporter 3′ RACE products were IPA transcripts under control conditions (49.3%, Fig. 2B, lane 4) and the remaining spliced RNAs were due to the usage of a cryptic 5′ ss (50.7%, Fig. 2B, lane 4, band marked by an arrow, and Fig. S2C). Additionally, U1 AMO treatment led to a nearly complete switch to IPA transcripts (Fig. 2B, lane 5). These data suggest that a weaker 5′ ss led to reduced splicing efficiency of a specific intron and higher IPA site usage. Similarly, when the 3′ ss was mutated to a weaker site, significant levels of IPA transcripts were observed (15%) while the spliced transcripts were due to the use of a cryptic 3′ ss (Fig. 2B, lane 6, band marked by an arrow, and Fig. S2C). U1 AMO treatment again caused the production of almost exclusively IPA transcripts (Fig. 2B, lane 7). When both the 5′ and the 3′ ss were mutated, IPA transcripts were produced predominantly and, conversely, spliced transcripts were detected predominantly when the IPA site was abolished (Fig. 2B, lanes 8-11). We also tested the effect of PB on these reporters and obtained very similar results to U1 AMO treatment (Fig. 2C), consistent with splicing-dependent repression of IPA. Thus, these results suggest that disrupting splicing of a specific intron by mutating either its 5′ or 3′ ss can lead to higher IPA site usage.

**Figure 2.**
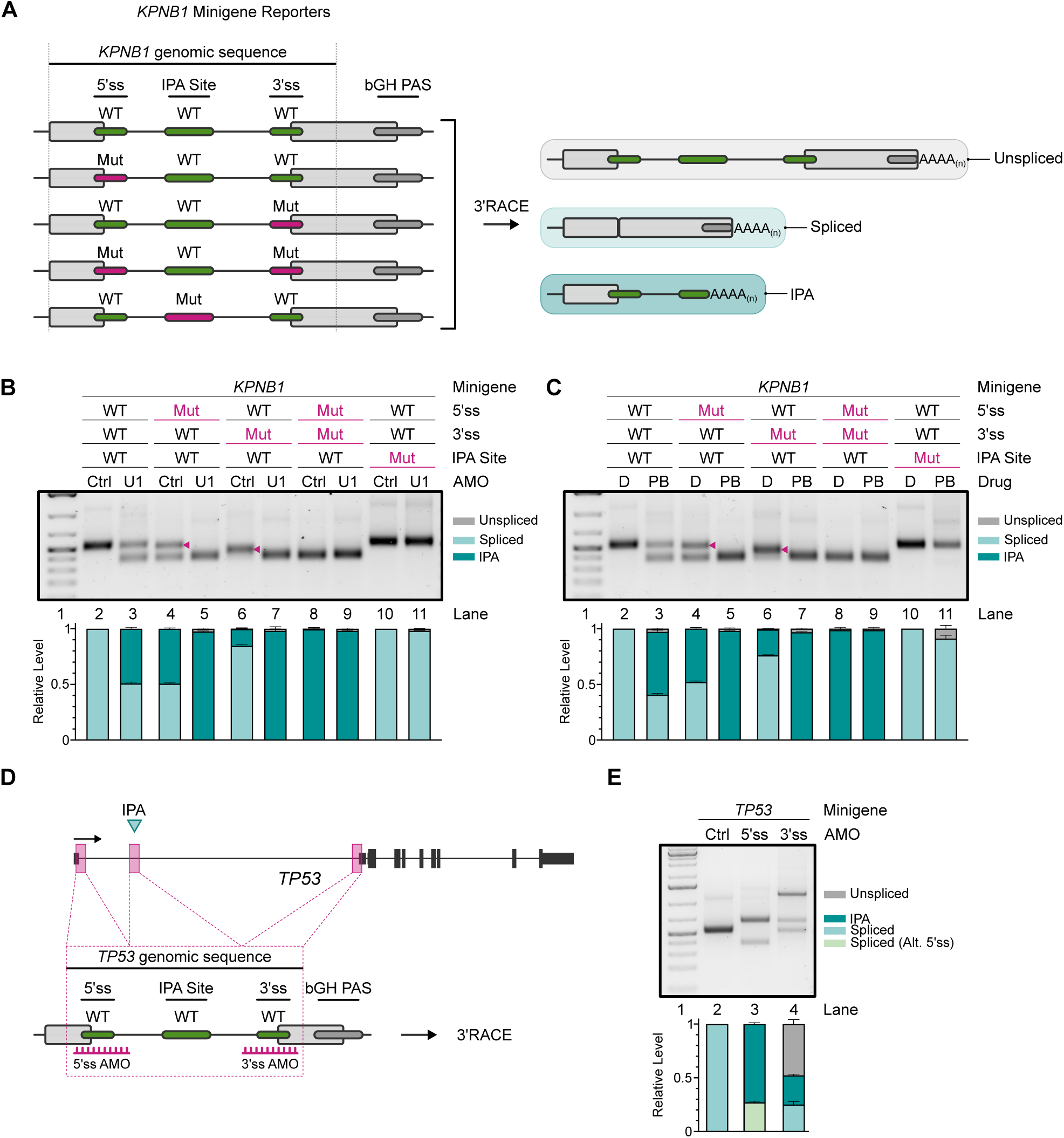
Gene-specific splicing inhibition activates intronic polyadenylation. **(A)** Left: Schematic of the *KPNB1* minigene reporters used in **B** & **C**. The reporters contain the genomic sequence of *KPNB1* spanning exons 13 and 14 and the intervening intron, which contains an IPA site. Green boxes indicate the original wild-type (WT) genomic sequences. Pink boxes indicate the mutated (Mut) splice sites or IPA site introduced by PCR mutagenesis. Right: Schematic of the 3′RACE products detected and quantified in **B** & **C**. **(B-C)** 3′ RACE analysis of the *KPNB1* minigene reporters in control-treated, U1 AMO-treated (**B**) or PB-treated (**C**) cells. Top: Agarose gels used to resolve 3′RACE products. Bottom: Quantification of 3′RACE performed using a fragment analyzer (see Methods). Each stacked barplot includes the relative abundance of unspliced, spliced, and IPA 3′RACE products in the specified sample. Data are presented as mean ± SD (*n* = 3). **(D)** Schematic of the *TP53* genomic structure (top) and sequences used to generate the *TP53* minigene reporter (bottom). The position of the splice-site specific AMOs is depicted in the schematic. **(E)** 3′ RACE analysis of the *TP53* minigene reporter in control-treated, 5′ss AMO-treated, or 3′ss AMO-treated cells. Top: Agarose gel used to resolve 3′RACE products. Bottom: Quantification of 3′RACE performed using a fragment analyzer (see Methods). Each stacked barplot includes the relative abundance of unspliced, spliced, spliced using an alternative 5′ ss, or IPA 3′ RACE products in the specified sample. Data are presented as mean ± SD (*n* = 3).

The vast majority of mammalian genes contain multiple exons and it has been proposed that internal exons are recognized by splicing factors such as U1 snRNP, U2 snRNP, and U2AF bound across the same exon through a mechanism called exon definition.^44^ Thus, disrupting one splicing factor could lead to a defect in exon definition, disrupting binding of other splicing factors, such as U1 snRNP, which could then indirectly de-repress IPA. To directly test this, we built a three-exon reporter by inserting the endogenous *KPNB1* upstream exon and intron into our two-exon reporter (Fig. S2G). We then mutated the 5′ or 3′ ss of the downstream intron to weaker sites, similar to Fig. 2B-C, and evaluated its impact on IPA. Our results showed that weakening the 5′ or 3′ ss of the downstream intron led to higher abundance of the IPA transcripts under control conditions, which was further increased by splicing inhibition by U1 AMO or PB treatment (Fig. S2H-I). Although we detected a small amount of splice site mutant reporter RNAs with an unspliced second intron upon U1 AMO or PB treatment, most of the transcripts under these conditions were IPA transcripts with the first intron fully spliced (Fig. S2H-I). This suggests that mutating the splice sites of the downstream intron did not significantly compromise splicing of the first intron, suggesting that the effect of splicing disruption on IPA activation was unlikely to be caused solely by a defect in exon definition. On the other hand, defective splicing of an intron could impact that splicing and IPA usage of a downstream intron, as discussed later.

In addition to blocking splicing by directly mutating splice site sequences, we also tested the effect of splicing inhibition by using intron-specific splice site targeting-AMOs. As shown in Fig. 1D, the tumor suppressor gene *TP53* contains an IPA site in its first intron, which was significantly induced by nearly all treatments that globally inhibited splicing, while the levels of the full-length transcript decreased. We designed a minigene reporter that expressed a shortened version of the genomic sequence spanning the first two exons of the gene *TP53*, which contains the IPA site (Fig. 2D). When we transfected the *TP53* reporter into control AMO-treated HEK293T cells, all of the detected reporter mRNAs were spliced (Fig. 2E, lane 2). However, when we co-transfected the reporter with an AMO that base-paired with the 5′ ss of the reporter (Fig. 2D), the majority (72.9%) of the *TP53* reporter mRNAs were IPA transcripts (Fig. 2E, lane 3), while the remaining transcripts were spliced using an alternative, cryptic 5′ ss (Fig. 2E, lane 3 and Fig. S2D). Similarly, co-transfecting an AMO that was complementary to the 3′ ss of the reporter intron led to a significant increase in the levels of IPA (27.2%) and unspliced (47.9%) mRNAs (Fig. 2E, lane 4). When the IPA site was mutated, 5′ and 3′ ss AMOs blocked splicing or induced splicing from a cryptic 5′ ss (Fig. S2E-F). Together our data demonstrated that inhibiting the splicing of a specific intron, by either mutating its 5′ or 3′ ss or by blocking them using AMOs, induced IPA site usage, suggesting that pre-mRNA splicing and CPA compete with each other during transcription.

### Intron sequence features modulate the specificity of splicing factor inhibition-induced IPA activation

Our results showed that splicing inhibition by targeting different splicing factors leads to activation of overlapping and distinct IPA sites (Fig. 1B-E and Fig. S1C-E). To investigate the molecular basis for such specificity, we first compared various features of IPA sites that were activated by specific splicing inhibitors and the introns in which these IPA sites reside, which will be herein referred to as host introns. Both PB and OTS964 target SF3b1 and their IPA activation profiles were highly similar (Fig. 1C). Likewise, U2AF1 and U2AF2 knockdown impacted a similar set of IPA events (Fig. 1C). For our analyses of the specificity underlying IPA activation, we combined IPA sites that were activated by PB with those that were activated by OTS964 (PB/OTS964) and, similarly, combined IPA sites that were activated by U2AF1 KD with U2AF2 KD-activated IPA sites (U2AF KD). Since there were no IPA sites that were only activated by U2 AMO, we did not include U2 AMO in our comparative analyses. By comparing the IPA sites activated by U1 AMO, PB/OTS964, and U2AF KD, we found that they targeted overlapping and distinct sets of IPA sites (Fig. 3A). Next, we characterized the position of the IPA host introns. For the following analyses, we also included constitutive IPA sites, which are defined as IPA sites that are used in 10% or more transcripts from a gene under control conditions. Although constitutive IPA sites were enriched in the last intron, IPA sites that were activated by all three groups (common), U1 AMO only, or U2AF KD only were highly enriched in the first intron (Fig. 3B). We next compared features of the IPA host introns and the IPA sites themselves and observed the following: 1) the upstream 5′ ss strength, as measured by the MaxEnt score,^45^ of constitutive IPA sites was significantly lower than other categories (Fig. 3C), consistent with our reporter assay results that weak splice sites lead to higher IPA site usage under normal conditions; 2) the host introns of U1 AMO- and U2AF KD-specific IPA sites were significantly larger than those all other groups (Fig. 3D); 3) U2AF KD-specific IPA sites were significantly stronger PAS, as measured by the APARENT score,^46^ than most groups (Fig. S3B), and were located significantly further from their upstream 5′ ss or downstream 3′ ss (Fig. S3C-D). These results suggest that the intron sequence context of IPA sites impact their response to specific splicing inhibitors.

**Figure 3.**
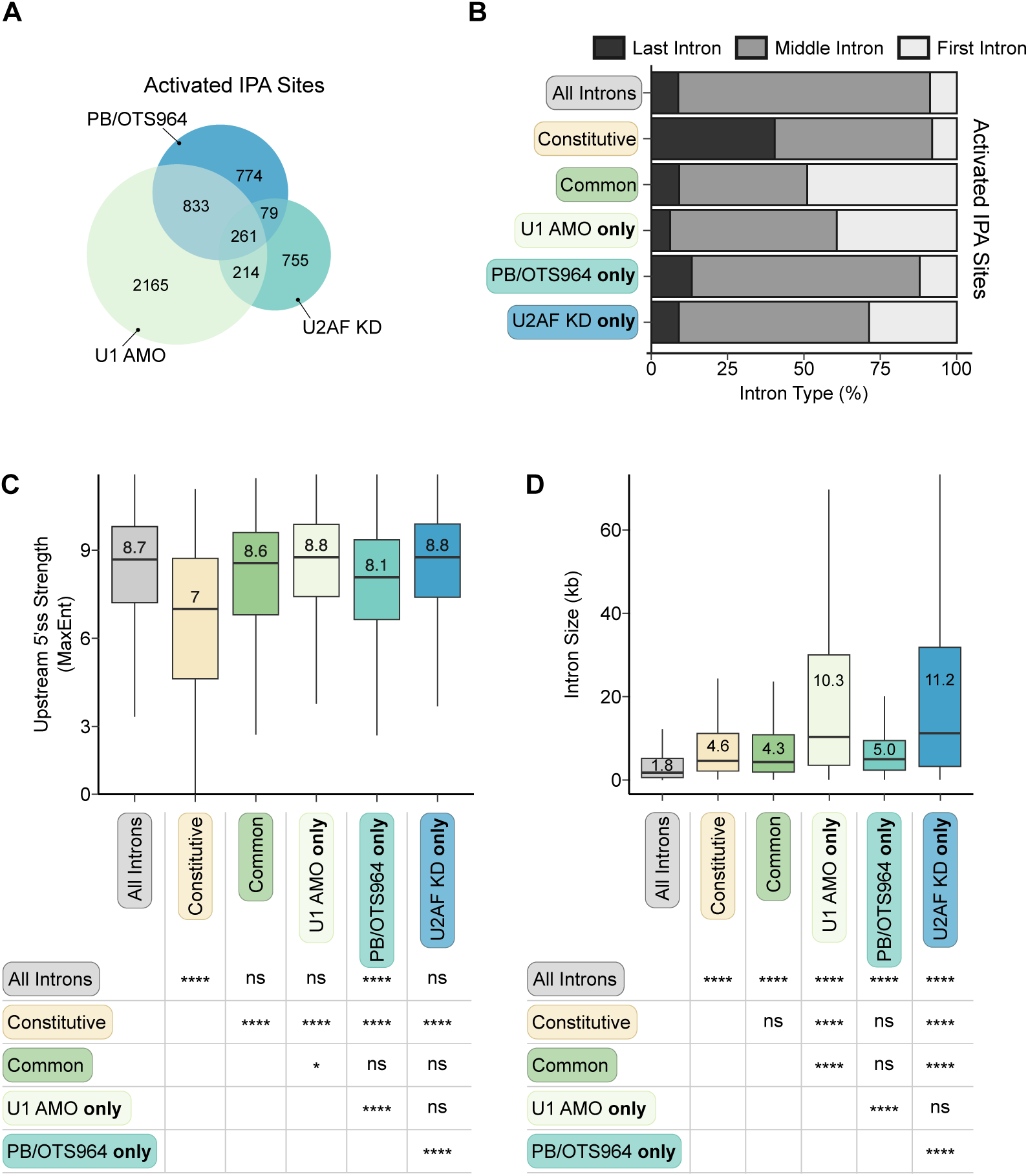
The intronic context of IPA sites impacts their response to specific splicing inhibitors. **(A)** Area proportional venn diagram of IPA sites activated by U1 AMO, PB/OTS964, and U2AF KD. Information from these IPA site groups is used in **B – D**. **(B)** Stacked barplot depicting the intron position of IPA sites that are constitutively used (Constitutive), activated by U1 AMO, PB/OTS, and U2AF KD (Common), or activated by U1 AMO only, PB/OTS964 only, or U2AF KD only. **(C)** Boxplots depicting the strength of the nearest upstream 5′ ss as measured by MaxEnt for IPA sites that are constitutively used (Constitutive), activated by U1 AMO, PB/OTS, and U2AF KD (Common), or activated by U1 AMO only, PB/OTS964 only, or U2AF KD only. Statistical analysis was calculated by Kruskal-Wallis test. ns p value > 0.05, ∗p value ≤ 0.05, ∗∗p value ≤ 0.01, ∗∗∗p value ≤ 0.001, ∗∗∗∗: p-value ≤ 0.0001. **(D)** Boxplots depicting the size of the intron that contains IPA sites that are constitutively used (Constitutive), activated by U1 AMO, PB/OTS, and U2AF KD (Common), or activated by U1 AMO only, PB/OTS964 only, or U2AF KD only. Statistical analysis was calculated by Kruskal-Wallis test. ns p value > 0.05, ∗p value ≤ 0.05, ∗∗p value ≤ 0.01, ∗∗∗p value ≤ 0.001, ∗∗∗∗: p-value ≤ 0.0001.

To further understand the molecular mechanism underlying the specificity of splicing inhibitor-induced IPA activation, we performed mRNA-seq analyses of all splicing inhibitor-treated cells and analyzed the intron retention (IR) levels for all introns as a proxy for splicing efficiency. Our results showed that all splicing inhibitors led to widespread increase in IR levels, but different splicing inhibitors compromised the splicing of overlapping and distinct sets of introns (Fig. 4A). Hierarchical clustering of IR levels showed that PB and OTS964 clustered together and U2AF1 and U2AF2 KD affected the splicing of a similar set of introns (Fig. 4A). This pattern was very similar to that of their IPA activation profiles (Fig. 1C). When we compared splicing inhibitor-induced IPA activation and intron retention patterns, we found that IPA activation by each splicing inhibitor was specifically and significantly associated with increased IR levels in the introns upstream of the IPA host intron (Fig. 4B-D and Fig. S4A). This pattern is illustrated in the example genes *TARDBP*, *TTI1*, and *HEXA* (Fig. 4E and Fig. S4B-C). In each case, there is significant IR in the intron immediately upstream of the IPA host intron in a splicing inhibitor-specific manner. This phenomenon, which we refer to as a “domino effect”, suggests that defective splicing of one intron could compromise splicing of a downstream intron, which may contribute to IPA site activation.

**Figure 4.**
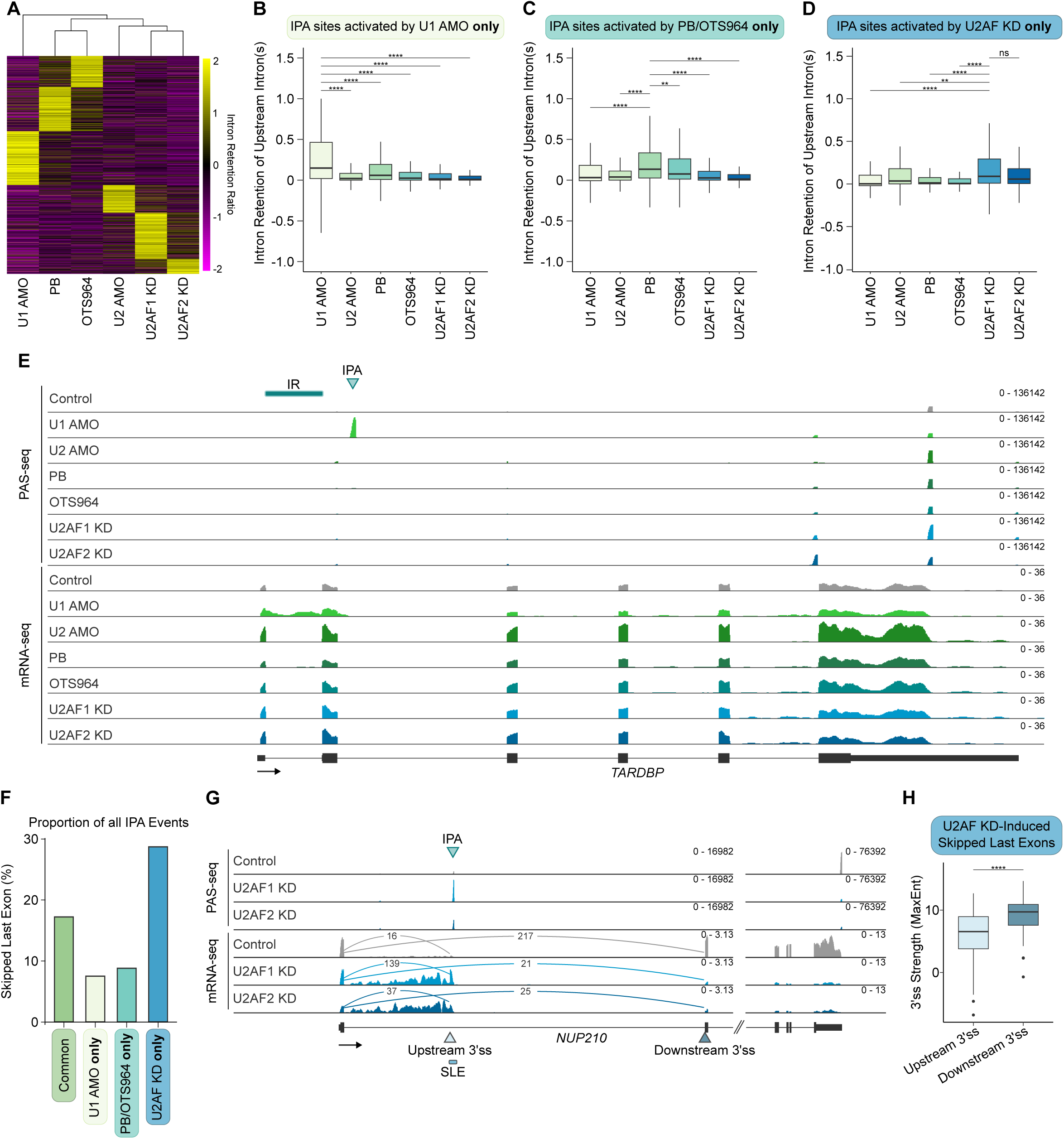
Splicing factor inhibition affects splicing and IPA in a coordinated manner. **(A)** Heatmap of the intron retention (IR) ratio scaled by row across all treatments. A positive value (yellow) indicates that the intron is retained more upon splicing inhibition. Hierarchical clustering was performed across treatments. **(B - D)** Boxplots depicting the IR ratio of introns upstream of IPA sites that are exclusively activated by U1 AMO **(B)**, PB/OTS964 **(C)**, or U2AF KD **(D)**. Statistical analysis was calculated by Kruskal-Wallis test. ns p value > 0.05, ∗p value ≤ 0.05, ∗∗p value ≤ 0.01, ∗∗∗p value ≤ 0.001, ∗∗∗∗: p-value ≤ 0.0001. **(E)** PAS-seq and mRNA-seq data tracks for the gene *TARDBP* following the indicated splicing inhibition treatments. Each peak in PAS-seq represents reads mapped to the 3′ end of mRNAs. IR is depicted by a bar and IPA is indicated by an arrow. **(F)** Barplot depicting the percentage of activated IPA events by each indicated treatment group that are skipped last exons. **(G)** PAS-seq and mRNA-seq data tracks for the gene *NUP210* following U2AF1 or U2AF2 KD. The activated cryptic 3′ ss (upstream 3′ ss), canonical downstream 3′ ss, activated skipped last exon (SLE), and IPA site are labeled. Sashimi plots are shown to indicate read counts that correspond to splicing to the upstream or downstream 3′ ss. **(H)** Boxplot comparing the 3′ ss strength of U2AF KD-activated cryptic upstream 3′ ss and downstream 3′ ss. Statistical analysis was calculated by t-test. ∗∗∗∗: p-value ≤ 0.0001.

In addition to increased IR, our mRNA-seq analyses also revealed that many splicing inhibitor-induced IPA sites were associated with skipped last exons (SLE), which are defined by a cryptic intronic 3′ ss and a downstream IPA site (Fig. S1B). Notably, 29% of IPA sites specifically induced by U2AF KD were associated with SLEs while this was true for only 7.5% and 8.8% of U1 AMO- and PB/OTS964-specific IPAs respectively (Fig. 4F). This effect is evident in the gene *NUP210*, in which U2AF KD led to increased splicing to a cryptic 3′ ss and IPA in the first intron (Fig. 4G). The activated 3′ ss of the SLE were significantly weaker than the corresponding downstream 3′ ss (Fig. 3H), which could explain why they were not used under normal conditions. Given that U2AF-specific IPA sites are significantly stronger than many other IPA sites (Fig. S3B), last exon definition mediated by these cryptic 3′ ss and IPA sites may outcompete splicing when U2AF is limited. Thus the preferential activation of SLE-associated IPA by U2AF is consistent with its essential function in 3′ ss recognition.^1^ Together our results suggest that splicing factors regulate splicing and IPA usage in a coordinated manner.

### U1 snRNP inhibition induces IPA-coupled and -independent premature transcription termination

Our analysis showed that, among all splicing inhibitors, IPA sites that were specifically induced by U1 AMO were highly enriched in the first intron (Fig. 3B). As CPA is tightly coupled to transcription,^47–49^ we next investigated whether the distribution of the IPA sites activated by different splicing inhibiting treatments was influenced by RNAPII transcription under these conditions. To this end, we performed 4-thiouridine labeled nascent RNA-sequencing (4sU-seq) of control cells or cells treated with each splicing inhibitor. To deduce transcription activities from the distribution of 4sU-labeled RNAs in these samples, we focused on the distribution of 4sU-seq signals along a composite intron that combines all introns of a gene, which correspond to RNAs in unprocessed pre-mRNAs. Strikingly, we observed dramatic accumulation of 4sU-labeled RNAs close to the start of the intronic region followed by a sharp decrease in U1 AMO-treated cells (Fig. 5A). This is consistent with a recent study showing that U1 inhibition leads to widespread premature transcription termination (PTT) in gene bodies.^36^ In contrast, the distribution of 4sU-labeled RNAs did not change significantly in cells treated with other splicing inhibitors (Fig. 5A). Thus, the accumulation of RNAs at the start of introns followed by PTT in U1 AMO-treated cells are likely to limit where IPA activation could occur, providing a potential explanation for the enrichment of U1 AMO-induced IPA sites in the first intron (Fig. 3B). For example, the majority of transcripts for both the *MED13* and *EFCAB14* genes were polyadenylated at poly(A) sites within their terminal exon in control cells and U1 AMO treatment caused a dramatic shift to an IPA site within an early intron (Fig. 5B-C). In both genes, PB treatment mostly activated a different IPA site, which was located downstream of the IPA site activated by U1 AMO (Fig. 5B-C). Similarly, the 4sU-seq signal in both genes was observed throughout the gene in control cells (Fig. 5B-C). By contrast, the 4sU-seq signal accumulated significantly in the early introns and decreased precipitously after the activated IPA sites, displaying hallmarks of premature transcription termination (Fig. 5B-C). Interestingly, U1 AMO-induced both IPA and PTT occurred upstream of those induced by PB treatment, suggesting that these two processes are connected. Together, these data suggest that the specificity of IPA activation by splicing inhibiting treatments is influenced by RNAP II transcription elongation and termination.

**Figure 5.**
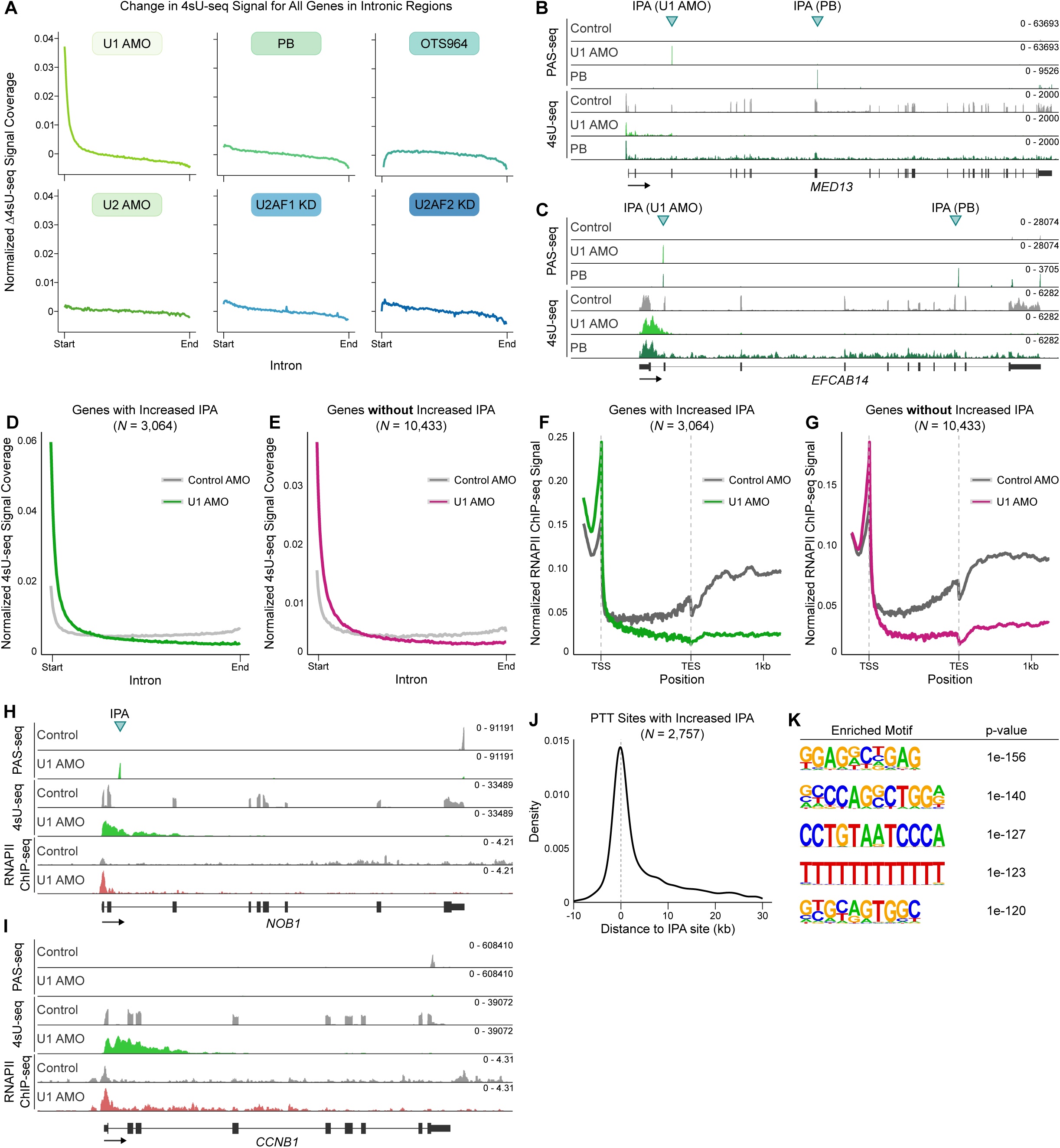
U1 snRNP inhibition induces IPA-coupled and IPA-independent premature transcription termination. **(A)** Metagene plots depicting the change in 4sU-seq signal for all genes following global splicing inhibition treatments. The x-axis depicts a composite intron in which all introns of each gene were combined to determine the relative position of 4sU-seq signal in all genes. **(B - C)** PAS-seq and 4sU-seq data tracks for the genes *MED13* (**B**) and *EFCAB14* (**C**) following global splicing inhibition by U1 AMO or PB. In PAS-seq, each peak represents reads mapped to the 3′ end of mRNAs. **(D)** Metagene plot depicting the 4sU-seq signal in control-treated cells (gray line) or U1 AMO-treated cells (green line) for all expressed genes that contained an activated IPA site following U1 AMO treatment as detected by PAS-seq (*N* = 3,064). The x-axis depicts a composite intron in which all introns of each gene were combined to determine the relative position of 4sU-seq signal. **(E)** Metagene plot depicting the 4sU-seq signal in control-treated cells (gray line) or U1 AMO-treated cells (pink line) for all expressed genes that did not contain an activated IPA site following U1 AMO treatment as detected by PAS-seq (*N* = 10,433). The x-axis depicts a composite intron in which all introns of each gene were combined to determine the relative position of 4sU-seq signal. **(F)** Metagene plot depicting the RNAPII ChIP-seq signal in control-treated cells (gray line) or U1 AMO-treated cells (green line) for all expressed genes that contained an activated IPA site following U1 AMO treatment as detected by PAS-seq (*N* = 3,064). The x-axis depicts a composite intron in which all introns of each gene were combined to determine the relative position of ChIP-seq signal. **(G)** Metagene plot depicting the RNAPII ChIP-seq signal in control-treated cells (gray line) or U1 AMO-treated cells (pink line) for all expressed genes that did not contain an activated IPA site following U1 AMO treatment as detected by PAS-seq (*N* = 10,433). The x-axis depicts a composite intron in which all introns of each gene were combined to determine the relative position of ChIP-seq signal. **(H - I)** PAS-seq, 4sU-seq, and RNAPII ChIP-seq data tracks for the genes *NOB1* (**H**) and *CCNB1* (**I**) following splicing inhibition by U1 AMO. In PAS-seq, each peak represents reads mapped to the 3′ end of mRNAs. *NOB1* (**H**) contains a U1 AMO-activated IPA site. *CCNB1* (**I**) does not contain any detected U1 AMO-activated IPA sites. **(J)** Density plot depicting the distribution of PTT site locations relative to the location of an U1 AMO-activated IPA site (*N* = 2,757). **(K)** Enriched motif analysis of the top 1000 PTT sites as detected by PTTseek.

Although U1 AMO induced IPA in over 3,000 genes, most genes did not display IPA activation. Interestingly, however, we observed a similar pattern of 4sU-labeled RNA signal in genes with and without IPA activation (Fig. 5D-E), suggesting that PTT also occurred in genes without IPA activation. To further validate that the 4sU-seq pattern reflects that of RNAPII occupancy, we analyzed published RNAPII ChIP-seq data in control or U1 AMO-treated cells.^50^ Indeed, we observed accumulation of RNAPII ChIP-seq signal near the transcription start sites (TSS) followed by a sharp decrease within gene bodies in genes with U1 AMO-activated IPA and those without (Fig. 5F-G). When we analyzed published RNAPII ChIP-seq from PB-treated cells, we observed evidence of more modest PTT in genes with or without PB-activated IPA (Fig. S5A-B).^51^ Such U1 AMO-induced PTT is evident in two example genes, *NOB1* and *CCNB1*, as evidenced by the 4sU-seq and RNAPII ChIP-seq data (Fig. 5H-I). However, IPA activation was observed only in *NOB1* but not in *CCNB1* (Fig. 5H-I). These results suggest that U1 inhibition leads to widespread PTT, which can occur in both an IPA-coupled and IPA-independent manner.

To begin to understand the scope and mechanism for U1 inhibition-induced PTT, we have carried out several analyses. First, by analyzing 4sU-seq data, we estimated that 61% of genes displayed U1 AMO-induced PTT (Fig. S5C-D, see Methods for details). The expression levels of genes with U1 AMO-induced PTT were significantly lower than those without PTT (Fig. S5E, p = 0.01, t-test). Second, the distribution of 4sU-seq signal in U1 AMO-treated cells—accumulation near the TSS followed by a sharp decline within the gene body—resembled that in cells depleted of the Integrator complex, which functions to attenuate transcription.^52^ To test if Integrator is required for U1 inhibition-induced PTT, we generated an Integrator degron cell line by fusing a FKBP12^F36V^ degron to Integrator 11 (INTS11) via genome editing.^53^ Immediately after treating cells with control or U1 AMO, we added either DMSO (vehicle) or the small molecule dTAG to deplete INTS11 (Fig. S5F) and then performed 4sU-seq analyses. Depletion of INTS11 in control AMO-treated cells led to accumulation of 4sU-labeled RNAs near the start of intronic regions (Fig. S5G), consistent with previous reports.^52^ In U1 AMO-treated cells, INTS11 depletion led to a modest extension of 4sU-seq signal within gene body, but PTT persisted under this condition (Fig. S5H), suggesting that the Integrator complex is not required for U1 AMO-induced PTT. Third, to investigate the mechanism underlying U1 AMO-induced PTT in an unbiased manner, we systematically identified the PTT sites in U1 AMO-treated cells using our 4sU-seq data and a new computational method, called PTTseek (see Methods) by detecting sharp declines in 4sU-seq signals. For genes with U1 AMO-induced IPA activation, the PTT sites tended to cluster near the IPA sites (Fig. 5J), suggesting that IPA activation and PTT are generally coupled in these genes. When we performed a motif analysis of all PTT sites, regardless of whether they were associated with IPA activation, we detected two major motifs, a G/C-rich motif and a T-tract motif (Fig. 5K). These sequences were reminiscent of those recently reported for U1 inhibition-induced “transcription transition sites”.^36^ Interestingly both of these motifs have been shown to be important for transcription termination in bacteria.^54^ Recent studies also suggested that the T-tract sequence promotes spontaneous transcription termination by RNAPII in eukaryotes.^55,56^ Together our results strongly suggest that U1 inhibition renders RNAPII transcription susceptible to PTT, either in an IPA-coupled or IPA-independent manner.

### Inhibition of pre-mRNA 3′ processing increases splicing efficiency

We showed that global and gene-specific splicing perturbations reduce splicing efficiency and increase IPA. This is consistent with a model in which splicing and IPA are competing RNA processing reactions. A prediction of this model is that reduced 3′ processing efficiency should increase splicing efficiency. To test this, we analyzed splicing and 3′ processing patterns following treatment with JTE-607, a small molecule inhibitor of the 3′ processing endonuclease CPSF73.^13,14,16^ Our previous PAS-seq analyses of DMSO- and JTE-607-treated cells showed that this inhibitor induced a widespread shift in PAS usage from proximal PAS to distal sites.^13^ When we focused our analysis on IPA transcripts, we found that JTE-607 broadly inhibited IPA site usage. Specifically, 151 genes showed significantly decreased IPA site usage (genes with change in PAS usage > 15% and reduced IPA usage, FDR < 0.05), whereas only 20 genes displayed increased IPA site usage (Fig. 6A). To evaluate the impact of CPA inhibition on splicing, we analyzed a published mRNA-seq dataset of DMSO- and JTE-607-treated cells^16^ and found that JTE-607 treatment led to significantly lower intron retention in 187 introns while the opposite change was only observed in 60 introns (change in intron retention > 15% and FDR < 0.05) (Fig. 6B), suggesting that CPA inhibition indeed stimulated splicing efficiency. Interestingly, however, the introns that displayed improved splicing efficiency following JTE-607 treatment generally lacked IPA sites (Fig. 6C), indicating that the improved splicing efficiency was not due to a change in IPA usage within the same intron. Therefore, CPA inhibition could improve splicing efficiency in at least two different ways. First, as shown for the *KPNB1* minigene reporter (Fig. 2A-C and Fig. S6), and the genes *BFAR* (Fig. 6D) and *DIDO1* (Fig. 6E), 3′ processing inhibition by mutating the PAS or treatment with JTE-607 resulted in a significant decrease in the IPA transcript levels and a concomitant increase in splicing of the intron in which the IPA site resides. Second, in JTE-607-treated cells, many poorly spliced introns (retained introns) displayed higher splicing efficiency, as evidenced by lower mRNA-seq signal within the intron in JTE-607-treated cells (Fig. 6F-G). One possible explanation for this observation is that less efficient 3′ processing may lead to increased dwell time in the nucleus for all transcripts, including those that contain unspliced introns. Such prolonged residence in the nucleus may allow more time for the splicing machinery to remove these introns. Together, our data provides strong evidence that splicing and 3′ processing are in competition with each other and the underlying mechanisms will be discussed below.

**Figure 6.**
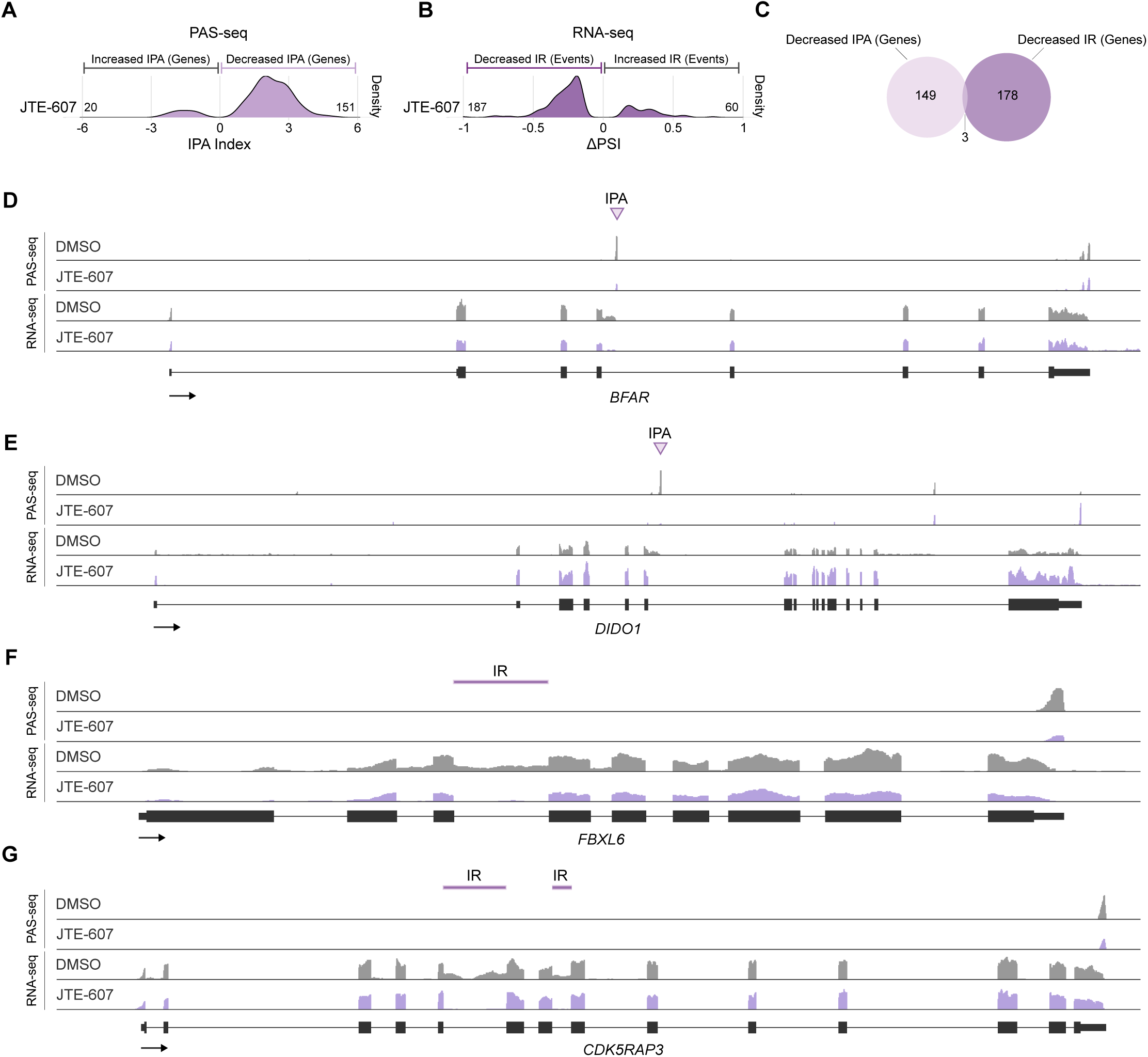
Inhibition of pre-mRNA 3′ processing increases splicing efficiency. **(A)** Density plot depicting the change in IPA site usage following treatment of cells with JTE-607. Values to the left of 0 indicate genes with increased IPA site usage. Values to the right of 0 indicate genes with decreased IPA site usage. The numbers of genes that contained significantly upregulated or downregulated IPA sites (genes with >15% change in PAS usage and increased (upregulated) IPA site usage or decreased (downregulated) IPA site usage, FDR < 0.05) are depicted for each treatment.

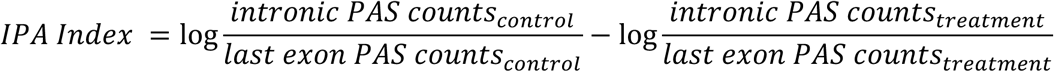 **(B)** Density plot depicting the change in percent spliced in (PSI) for retained introns following treatment of cells with JTE-607 as detected from mRNA-seq data using rMATS-turbo. Values to the left of 0 indicate introns with decreased intron retention. Values to the right of 0 indicate introns with increased intron retention. The numbers of introns that displayed significantly increased or decreased intron retention (introns with >15% change in intron retention, FDR < 0.05) are indicated. **(C)** Area proportional venn diagram of genes with decreased IPA as analyzed in **A** and decreased intron retention as analyzed in **B**. **(D**-**G)** PAS-seq and mRNA-seq data tracks for the genes *BFAR* (**D**), *DIDO1* (**E**), *FBXL6* (**F**), and *CDK5RAP3* (**G**) following 3′ processing inhibition by JTE-607. In PAS-seq, each peak represents reads mapped to the 3′ end of mRNAs. *BFAR* (**D**) and *DIDO1* (**E**) contain an IPA site that decreases in usage upon JTE-607 treatment but did not exhibit decreased intron retention upon JTE-607 treatment. *FBXL6* (**F**) and *CDK5RAP3* (**G**) contain introns that displayed decreased intron retention upon JTE-607 treatment but no IPA was detected in these introns in control- or JTE-607-treated cells by PAS-seq.

## DISCUSSION

It remains an important outstanding question how different pre-mRNA processing steps are coordinated with one another and with transcription to ensure efficient and accurate gene expression. Here we address this question by characterizing the relationship between splicing and 3′ processing. The prevailing telescripting model posits that U1 snRNP inhibits 3′ processing via a splicing-independent mechanism. This model was based on global analyses of U1 AMO-treated cells and more limited characterization of other splicing inhibitors.^31,32^ Here we inhibited splicing by using six different methods and showed that all tested splicing-inhibiting treatments activated 3′ processing at IPA sites. These treatments included AMOs, small molecule inhibitors, and RNAi depletion that targeted U1 and U2 snRNP, U2AF, SF3b1, and CDK11. This is consistent with recent studies showing that SSA and PB, both small molecule inhibitors of SF3b, and an AMO targeting U4 snRNA broadly activated IPA.^33–35^ Conversely, we demonstrated that inhibiting 3′ processing by using a small molecule JTE-607 promoted splicing efficiency. This is in keeping with a recent report showing that depletion of CPSF73, the endonuclease for 3′ processing and target of JTE-607, led to improved splicing.^33^ Together these results strongly suggest that splicing and 3′ processing are competing processing steps and that the balance between them modulates the outcome of gene expression.

Among all splicing-inhibiting treatments that we tested, the AMO that targeted U1 snRNA activated a greater number of IPA sites than other treatments (Fig. 1B and Fig. S1C). The different potency in IPA activation among splicing inhibiting treatments could be linked to the functions of these splicing factors and the mechanism of action for these splicing-inhibiting treatments. Although U1 snRNP is believed to be required for recognizing all 5′ ss, U2AF is only involved in recognizing a subset of 3′ ss. For example, for introns with long stretch of polypyrimidine tracts, U2AF1 is dispensable for 3′ ss recognition.^57^ Additionally, a subset of short introns rely on SPF45/RBM17, but not U2AF, for 3′ ss binding.^58^ Similarly, small molecule inhibitors of SF3b1 have been shown to block splicing in a sequence-dependent manner.^59,60^ This is consistent with our observation that blocking different splicing factors compromised the splicing of overlapping and distinct sets of introns (Fig. 4A). Despite the differences in the potency and selectivity of these splicing-inhibiting treatments, the fact that blocking ss or branch point recognition (U1 and U2 snRNPs, U2AF, SF3b1) (this study and ^31,33,34^) and splicing catalysis (U4 snRNP)^35^ all lead to IPA activation strongly suggest a competitive relationship between splicing and 3′ processing.

It remains unclear how splicing and 3′ processing compete with each other. Given the co-transcriptional nature of many splicing and CPA events, there is kinetic competition between these processes.^61,62^ On the other hand, their competition likely extends beyond the mere difference in the speed of these two processes. Recent studies suggest that the 3′ processing cleavage reaction occurs quickly, with half-lives under one minute.^63^ As many IPA sites are located within large introns and far from the 3′ ss (e.g. Fig. 1D-E and Fig. S1D-E), one would predict that 3′ processing could be completed before the 3′ ss is transcribed. Thus 3′ processing within introns is most likely actively suppressed by splicing. Previous studies proposed that U1 snRNP is tethered to RNAPII during transcription to allow splicing to take place quickly after the 3′ ss is transcribed.^64,65^ Consistent with this model, recent structural studies showed that the 5′ ss-bound U1 snRNP can bind to the core of RNAPII.^66^ On the other hand, previous studies provided evidence that CPA factors also bind to the core of RNAPII for co-transcriptional 3′ processing.^67^ As such, there may be a competition between splicing and CPA factors for binding to RNAPII during transcription. When RNAPII transcribes through introns, U1 snRNP and perhaps other splicing factors preferentially bind to RNAPII and block access for CPA factors, thereby inhibiting intronic 3′ processing. Consistent with this model, earlier studies have shown that U1 AMO not only blocks U1 snRNA-5′ ss base-pairing, but also dissociates U1 snRNP from RNAPII.^68^ Depletion of other elongation factors, such as the PAF complex and SCAF4/8, has been shown to lead to premature transcription termination and 3′ processing at IPA sites.^69–71^ Additionally, previous studies provided extensive evidence that splicing and CPA influence each other via direct interactions among splicing and CPA factors.^24,72–75^ Thus, the decision between splicing and 3′ processing depends on a complex interplay among transcription, splicing and CPA machineries. Furthermore, the “domino effect” we observed, i.e. defective splicing of an upstream intron affecting the splicing and IPA of a downstream intron (Fig. 4B-E), suggest that the competition may occur beyond individual introns.

A recent study^36^ and our results demonstrated that inhibiting U1 snRNP leads to widespread PTT within gene body. Although some PTT events are accompanied by IPA activation, the majority seem to occur without cleavage/polyadenylation. The Integrator complex has been shown to promote RNAPII premature termination, but out data suggest that Integrator does not play a major role in U1 AMO-induced PTT. Our analysis of the PTT sites detected two major motifs, a G/C-rich motif and a T-tract. Both of these motifs have been shown to be important for transcription termination in bacteria^54^ and recent studies have reported that the T-tract sequence promotes spontaneous transcription termination by RNAPII in eukaryotes.^55,56^ These data indicate that the fundamental mechanism for transcription termination may be more conserved between prokaryotes and eukaryotes than previously thought, and that the emergence of splicing may have provided an additional layer of regulation on transcription in eukaryotes. Interestingly, a recent study showed that U1 snRNP and the elongation factor SPT6 cooperatively bind to RNAPII.^76^ This interaction was proposed to stimulate spliceosome assembly, but it is likely that the interactions between U1 snRNP and elongation factors also stimulate transcription elongation and/or prevent termination.

At least 30% of annotated 3′ processing sites are within introns.^27^ IPA transcripts often do not encode proteins or rather encode truncated non-functional proteins. Thus, IPA activation can be functionally equivalent to gene inactivation. Indeed, it has been proposed that IPA activation leads to inactivation of many large genes, including tumor suppressor genes.^28,30^ Additionally, the noncoding IPA transcript from the *TP53* gene has recently been shown to be oncogenic.^29^ A number of small molecular inhibitors of splicing, such as PB and its derivatives, Spliceostatin A, and OTS964, are considered as potential therapeutic reagents.^11,17–20^ However, as treating cells with these molecules led to IPA activation in many genes, including tumor suppressor genes such as *TP53* and *BECN1* (Fig. 1D-E), they may have harmful consequences. Thus, our study suggests that caution must be taken when using general splicing inhibitors to treat diseases.

## Data Availability

PAS-seq datasets have been deposited into the GEO under accession number GSE275154.

## Code Availability

The python script PTTseek.py has been deposited into GitHub: https://github.com/yongshes/PTTSeek

## Acknowledgments

We would like to thank Drs. Klemens Hertel, Shalini Sharma, and Doug Black for providing reagents, and Feng Qiao for comments on the manuscript. We wish to acknowledge the support of the Chao Family Comprehensive Cancer Center Shared Resource Genomics High-Throughput Facility, supported by the National Cancer Institute of the National Institutes of Health under award number P30CA062203. We would like to thank Seung-Ah Yoon from the UCI Genomics High-Throughput Facility, and Adeela Syed from the UCI Optical Biology Core Facility for technical support. This study was supported by the following grants: R35GM149294 (Y.S.). L.T. is supported by the Hewitt Foundation Postdoctoral Fellowship. M.C.V is supported by a NIH Diversity Supplement (R01AI170840-03S1).

## Author Contributions

Conceptualization: L.V.S. and Y.S.; Investigation: L.V.S., S.L., L.L., K.S.K., Y.Y.; Formal Analysis: L.V.S., E.A., I.M., Y.S.; Validation: L.V.S., S.L., M.C.V., L.T., L.L.; Writing – Original Draft: L.V.S.; Writing – Review & Editing: L.V.S., Y.S.; Funding Acquisition: Y.S.; Supervision: Y.S.

## STAR★ METHODS

## EXPERIMENTAL MODEL AND STUDY PARTICIPANT DETAILS

### Cell Lines

Wild-type HEK293T, edited HEK293T, and Flp-in™ T-REx™-293 cell lines were maintained in DMEM (Gibco) supplemented with 10% FBS (Sigma). Cells were grown at 37°C with 5% CO_2_. Cell culture media for Flp-in™ T-REx™-293 parental cells was additionally supplemented with 15 µg/mL blasticidin and 100 µg/mL zeocin. Cell culture media for Flp-in™ T-REx™-293 stably edited cell lines was supplemented with 15 µg/mL blasticidin and 500 µg/mL G418. Cell lines were routinely monitored for mycoplasma contamination using the MycoAlert Mycoplasma Detection Kit (Lonza).

## METHOD DETAILS

### Splicing Inhibition Treatments

#### Inhibition of U1 snRNP or U2 snRNP by AMO

To inhibit U1 snRNP, 25 µM control or U1 AMO (GeneTools, LLC) was delivered into 1x10^6^ cells per cuvette by nucleofection using the Lonza SF Cell Line 4D-Nucleofector^TM^ X Kit using the DH-135 program. To inhibit U2 snRNP, 50 µM control or U2 AMO (GeneTools, LLC) was nucleofected following the same procedure. For the combined treatment of U1 AMO and INTS11 depletion, 25 µM control or U1 AMO was delivered into a total of 6x10^6^ cells per sample by nucleofection. Following nucleofection, cells were plated into a 10-cm plate containing 10 mL final media volume with a final concentration of 500 nM dTAG^v^-1 or an equivalent volume of DMSO. For all analyses, cells were harvested 16 hours later by direct lysis in Trizol.

#### Splicing Inhibition by Pladienolide B

To inhibit splicing via Pladienolide B (PB), PB or an equivalent volume of DMSO was added directly to cell culture media at a final concentration of 100 nM. For all analyses, cells were harvested 8 hours later by direct lysis in Trizol.

#### Splicing Inhibition by OTS964

To inhibit splicing via OTS964, OTS964 or an equivalent volume of DMSO was added directly to cell culture media at a final concentration of 50 nM. Cells were harvested 8 hours later by direct lysis in Trizol.

#### Depletion of U2AF1 and U2AF2 by Lentiviral Knockdown

Lentiviral particles were prepared by co-transfecting HEK293T cells with the following plasmids: pLKO: shRNA vector, PMD2.G, and psPAX2. The virus-containing cell culture media was collected 24 and 48 hours after transfection and passed through a 45 µM filter. The titer was measured using a qPCR-based kit (Applied Biological Materials). HEK293T cells were transduced at an MOI of 10 with lentivirus expressing one shRNA targeting U2AF1, two different shRNAs targeting U2AF2, or an empty vector. Polybrene (8 µg/mL) was included during transduction. 24 hours after transduction, puromycin was added to the media at a concentration of 1.25 µg/mL. 5 days after transduction, cells were harvested in Trizol for RNA purification or directly lysed in 1X SDS loading dye for western blotting.

### Generation of INTS11 FLAG-dTAG Edited HEK293T Cell Line

The N-terminal FLAG-dTAG knockin of INTS11 in HEK293T cells was achieved using CRISPR/Cas9 and the microhomology end joining (MMEJ) approach as described in,^77^ except that we used Lipofectamine 3000 to perform the transfections and used blasticidin (10 µg/mL) during antibiotic selection. To construct the donor vector, we modified pCRIS-PITCHv2-BSD-dTAG (BRD4) to replace the HA tag with a 3X-FLAG tag and replaced the microhomology sequences with those from INTS11 using In Fusion cloning (Takara). pCRIS-PITCHv2-BSD-dTAG (BRD4) were gifts from James Bradner & Behnam Nabet (Addgene plasmid # 91795 ; https://www.addgene.org/91795/ and Addgene plasmid # 91792 ; https://www.addgene.org/91792/).^78^ We used CHOPCHOP^79^ to select a specific and efficient sgRNA. We screened and validated homozygous edited single colonies using genomic DNA PCR followed by Sanger Sequencing by western blotting.

### PAS-seq

#### Library Preparation

PAS-seq libraries were prepared as previously described with the following minimal modifications.^40^ Purified mRNA was fragmented at 94°C for 3 minutes using the NEBNext Magnesium RNA fragmentation module. In addition, the libraries were resolved on a 2.5% agarose gel and the region between 200-300 base pairs was gel extracted and sequenced on the Illumina NovaSeq6000 platform. For all samples except U2AF1 KD and U2AF2 KD, a small amount of mouse RNA was spiked in but was not used in data processing. For the spike-ins, an extra control or treatment well was prepared and the cells in this extra well were counted and recorded. 1 µg total RNA from each sample was mixed with up to 100 ng (10%) spike-in mouse RNA (purified from mouse brain tissue) and used during library preparation. The spike-in RNA mass was calculated relative to the measured cell count for each sample. For example, if there were 1 x 10^6^ control cells and 0.8 x 10^6^ treated cells, then 100 ng mouse spike-in RNA would be added to 1 µg of control total RNA and 80 ng mouse spike-in RNA would be added to 1 µg of treated total RNA. As such, the level of spike-in RNA could be used to normalize sequencing reads to the original cell number.

#### PAS-seq Data Processing

Alternative polyadenylation analysis was conducted as previously described with few modifications.^80^ Briefly, we used Cutadapt (v2.10)^81^ to produce trimmed PAS-seq reads by: 1) trimming the 6 nucleotide linker sequence, 2) trimming the poly(A) tail sequence, and 3) removing any untrimmed reads. Trimmed reads were then mapped to a concatenated hg19 and mm9 genome using STAR (v2.7.3a)^82^ with the --alignEndsType EndToEnd parameter. The mapped reads were converted from a bam file to a bed file using bedtools (v2.29.2)^83^. Using a custom python script, reads were removed as potential internal priming events if they mapped to a genomic region where 6 consecutive A’s or 7 A’s out of 10 nucleotides were observed in the 10 nucleotides downstream of the read. To generate bigwig files, the remaining reads were first intersected using bedtools intersect with a master file of all annotated poly(A) sites, converted to a bam file, and bigwig files were generated using bedtools. The 3’ ends of the reads after internal priming removal were extracted from the bed files using bedtools flank and the initial read counts for each poly(A) site were calculated using bedtools coverage with a master file of all annotated poly(A) sites. Samples treated with pLKO, U2AF1 KD, or U2AF2 KD were processed the same as above except that they did not contain spiked in mouse RNA and thus reads were mapped to an hg19 genome. As the PAS-seq libraries prepared from JTE-607 treated HepG2 cells from a previously published study^13^ were prepared using a previous PAS-seq protocol, the PAS-seq data was processed the same as above, except that the removed nucleotide linker sequence was 4 nucleotides, rather than 6.

#### Alternative Polyadenylation Analysis

Poly(A) sites with significantly changed usage in the experimental condition were identified using diffSpliceDGE and topSpliceDGE within the edgeR package where each poly(A) site was treated as an “exon” (v3.40.1).^84^ To identify all affected genes and classify their primary change in alternative polyadenylation, we selected all genes with at least one poly(A) site that exhibited at least a 15% change in usage (increased or decreased) and met our cutoff for significance (FDR < 0.05). We then selected the two most differentially used poly(A) sites within each gene and calculated the difference in the proximal-to-distal ratio between control cells and treated cells. Of the two poly(A) sites, if the more proximal site was located in an intron and the more distal site was located in the last exon, this was classified as an IPA event. All other alternative polyadenylation changes were classified as non-IPA events.

#### Identification of Skipped Last Exons (SLE)

The introns detected in mRNA-seq mapping results were compared with detected the IPA sites in PAS-seq. If an IPA site was located between two 3′ ss with the same upstream 5′ ss, the region between the upstream 3′ ss and IPA were designated as a SLE.

#### KEGG Pathway Analyses

The Gene Ontology (GO) enrichment analysis and KEGG pathway analysis were performed by the Database for Annotation, Visualization and Integrated Discovery (DAVID) (https://david.ncifcrf.gov/home.jsp).^85,86^ The terms related to cell development, growth, and disease were collected. The terms in which p value < 0.05 and FDR < 0.05 were regarded as a significant functional annotation clustering.

### mRNA-seq and 4sU-seq

#### Sample Preparation

For all samples used for mRNA-seq and 4sU-seq, freshly prepared 4-thiouridine (4sU) (Sigma) was added directly to cell culture medium at a final concentration of 500 µM for the last 30 minutes of every splicing inhibition treatment or control condition. Cells were directly lysed in Trizol and total RNA was purified following the manufacturer’s protocol. For mRNA-seq, at least 500 ng total RNA was DNase-treated (Promega) and purified by phenol/chloroform extraction followed by ethanol precipitation and then directly used for library preparation.

#### Library Preparation

mRNA-seq Libraries were prepared using the Illumina TruSeq mRNA stranded kit with oligo dT-selection. For 4sU-seq only, total RNA was enriched for 4sU-labeled RNAs as previously described.^87^ 4sU-labeled RNAs were then used for library construction using the Illumina TruSeq Total RNA kit with ribosomal RNA depletion.

#### mRNA-seq and 4sU-seq Data Processing

Adaptor sequences and low-quality nucleotides were trimmed using trimGalore.^88^ Trimmed reads were then mapped to the hg19 genome using STAR (v2.7.3a)^82^ with the--alignEndsType Local parameter. The resulting bam files were used to generate bigwig files for visualization using bedtools (v2.29.2).^83^

#### Alternative Splicing Analysis

Alternative splicing analysis was performed on splicing inhibitor-treated cells and by analyzing a previously published a RNA-seq dataset prepared from whole cell lysate of A-673 cells treated with 10 µM JTE-607 or an equivalent volume of DMSO (NCBI Sequence Read Archive Accession: SRP158650).^16^ For JTE-607-treated samples, RNA-seq bam files produced after alignment were analyzed for changes in alternative splicing patterns using rMATS-turbo.^89^ The inclusion level difference (ΔPSI) was multiplied by -1 in order to depict decreases in intron retention (IR) following JTE-607 treatment as a negative value. To identify all significantly changed IR events, we filtered the RI.MATS.JC output for all IR events that met the following conditions: absolute value of ΔPSI ≥ 0.15 and FDR ≤ 0.05. For intron retention analysis of the splicing inhibitor-treated cells, intron retention level was calculated using IRFinder.^90^

#### Premature Transcription Termination Analyses

The distribution of 4sU-seq signal in intronic regions was analyzed by using the computeMatrix tools in the deepTools package.^91^ To analyze the composite introns from entire genes, we used the metagene option and a custom made hg19 intron bed12 file. Premature transcription termination (PTT) sites were detected in introns by using a custom script PTTseek.py using bigwig files of the 4sU-seq data and annotated intron bed file. To identify genes with PTT, 4sU-seq reads in each intron was counted using Rsubread for Control and U1 AMO-treated cells and analyzed using edgeR Splicing mode. Gene expression levels were quantified by using Salmon^92^ and those with TPM > 5 were selected. Genes whose read counts in the first intron significantly increased (FDR < 0.05 and logFC > 0) and those in the last intron significantly decreased (FDR < 0.05 and logFC < 0) were considered as having PTT.

### RNAPII ChIP-seq

#### Data Analysis

Previously published RNAPII ChIP-seq data was analyzed from HeLa cells treated with control or 15 nmol U1 AMO^50^ and K562 cells treated with DMSO or 1 µM PB.^51^ The RNAPII ChIP-seq signals were normalized with those from IgG and the meta-analysis of the distribution of the normalized signals along genes were performed using computematrix-region in deepTools.^91^

### Molecular Cloning

#### KPNB1 Minigene Reporters

The two-exon *KPNB1* minigene reporters included a 544-nucleotide sequence spanning the entire genomic region from the start of exon 13 to the end of exon 14. This sequence was amplified by PCR from genomic DNA and cloned into pCDNA5/ FRT/TO using In Fusion cloning. In the 5′ ss mutant, the 5′ ss sequence was mutated from 5′-GAG|GTGAGA-3′ to 5′-GTC|CATTCA-3′. In the 3′ ss mutant, the 3′ ss sequence was mutated from 5′-TTTTCTTATTTTCTTTGTAG|TCA-3′ to 5′-AAAAGAAAAAAAGAAAGATC|TCA -3′. In the PAS mutant, the intronic poly(A) signal was mutated from 5′-AATAAA-3′ to 5′-AAGAAA-3′. The three-exon *KPNB1* minigene reporter was created by inserting the genomic sequence for *KPNB1* exon 12 and intron 12 immediately upstream of the 2-exon reporter. All mutations were made by PCR mutagenesis.

#### TP53 Minigene Reporters

The *TP53* minigene reporters included a shortened version of the genomic sequence spanning from exon 1 to exon 2 of the *TP53* gene. As depicted in Fig. 2D, the *TP53* minigene included: the last 114 nucleotides of exon 1 (5′ UTR region) and 250 nucleotides of the immediate downstream intronic sequence to capture the entire 5′ ss, a 381 nucleotide sequence flanking the intronic poly(A) signal to capture the entire poly(A) site, and 250 nucleotides of intronic sequence upstream of exon 2 and the first 74 nucleotides of exon 2 to capture the entire 3′ ss. These fragments were amplified from genomic DNA and cloned into pCDNA5/FRT/TO using In Fusion cloning. In both the WT and Mut *TP53* minigene reporters, a cryptic 5′ ss sequence was mutated in exon 1 from 5′-CAG|GTAGCT-3′ to 5′-ACT|AGCGCC-3′ to prevent usage of this site following treatment with the 5′ ss-targeting antisense morpholino (AMO). In the *TP53* PAS mutant minigene reporter used in Fig. S2E, the canonical intronic poly(A) signal was mutated from 5′-AATAAA-3′ to 5′-AACAAA-3′ and a cryptic poly(A) signal located 5 nucleotides downstream of the canonical poly(A) signal was mutated from 5′-TATAAA-3′ to 5′-TACAAA-3′ to prevent usage of both sites in the PAS Mut control condition.

### Generation of *KPNB1* Minigene Reporter Stably Edited Cell Lines

To generate the *KPNB1* minigene reporter stably edited cell lines, 3x10^6^ Flp-in™ T-REx™-293 parental cells were plated in each well of a 6-well plate without any antibiotics in the media. The next day, 2.25 µg pOG44 and 250 ng pCDNA5 containing the desired *KPNB1* minigene were co-transfected using Lipofectamine 3000. The day after transfection, the media was changed to DMEM with 10% FBS and 15 µg/mL blasticidin. Two days after transfection, the cells were passaged to a 10-cm plate in DMEM with 10% FBS, 15 µg/mL blasticidin, and 500 µg/mL G418. Transfected cells then underwent polyclonal selection using 15 µg/mL blasticidin and 500 µg/mL G418 for approximately 14 days, or until a plate of untransfected control cells died.

### 3′ RACE of *KPNB1* and *TP53* Minigene Reporters

#### Expression of KPNB1 Minigene Reporters and Treatment with U1 AMO or Pladienolide B

To inhibit U1 snRNP, 25 µM U1 AMO or 25 µM Control AMO was nucleofected as described above into the Flp-in™ T-REx™-293 *KPNB1* reporter cell lines. Immediately after nucleofection, doxycycline was added to the cell culture media at a final concentration of 1 µg/mL. 16 hours later, the cells were lysed directly in Trizol. To inhibit splicing via PB, PB or an equivalent volume of DMSO was added directly to cell culture media at a final concentration of 100 nM. Immediately after addition of PB, doxycycline was added to the cell culture media at a final concentration of 1 µg/mL. 8 hours later, the cells were lysed directly in Trizol.

#### Expression of TP53 Minigene Reporters and Treatment with Splice Site-Targeting AMOs

The day before transfection, 50,000 HEK293T cells per well were seeded into a 24-well plate. The next morning, the media was removed and replaced with 500 µL of DMEM without FBS. 250 ng of pCDNA5: *TP53* minigene was then transfected using Lipofectamine 3000. Immediately after transfection, splice site-targeting AMOs (Vivo-Morpholinos, GeneTools LLC) were added directly to the media at the following final concentrations: Control AMO: 2.5 µM or 10 µM, 5′ ss AMO: 2.5 µM, 3′ ss AMO: 10 µM. 24 hours later, the cells were lysed directly in Trizol.

#### Expression of KPNB1 5′ss Mut Minigene Reporters and Treatment with JTE-607

To inhibit CPSF73 via JTE-607, JTE-607 or an equivalent volume of water was added directly to cell culture media at a final concentration of 10 µM. Immediately after addition of JTE-607, doxycycline was added to the cell culture media at a final concentration of 1 µg/mL. 8 hours later, the cells were lysed directly in Trizol and 3′ RACE was performed.

#### 3′ RACE

Total RNA was purified using Trizol following the manufacturer’s protocol. 500 ng or 1 µg total RNA was reverse transcribed using SuperScript III (ThermoFisher) with a custom 3′ RACE RT primer. The resulting cDNA was diluted by adding 40 µL water to the 20 µL RT reaction. Two consecutive PCR reactions were then performed to amplify the cDNA and monitor the splicing and intronic polyadenylation patterns. In the first PCR (PCR1), the forward primer (3′ RACE PCR1 F) was complementary to a region upstream of the first minigene exon and the reverse primer (3′ RACE PCR1 R) was complementary to the linker on the RT primer. PCR1 was performed using 1 µL of cDNA as the template and included 21 PCR cycles. In the second RACE PCR (PCR2), the forward primer (3′ RACE PCR2 F) was complementary to the region upstream of the first minigene exon, but downstream of the first forward primer binding site. The reverse primer was complementary to the linker on the RT primer (3′ RACE PCR2 R). PCR2 was performed using 1 µL of the PCR1 reaction as the template and consisted of 25 cycles. An aliquot of the PCR2 reaction was resolved on a 1% agarose gel and imaged. The remaining PCR2 reaction was purified using AmpureXP beads following the manufacturer’s protocol. An aliquot of the purified PCR2 reaction was analyzed using the DNF-473-0500 kit on the Agilent Fragment Analyzer. The relative quantities of unspliced, spliced, and intronically polyadenylated 3′ RACE products were quantified using the Agilent ProSize Data Analysis software.

### Primers used in this study

3′RACE RT primer: 5′-CCAGTGAGCAGAGTGACGAGGACTCGAGCTCAAGCTTTTTTTTTTTTTTTTTTTTV-3′

3′ RACE PCR1 F: 5′-CCATCCACGCTGTTTTGAC-3′

3′ RACE PCR1 R: 5′-CCAGTGAGCAGAGTGACG-3′

3′ RACE PCR2 F: 5′-AGCCTCCGGACTCTAGC-3′

3′ RACE PCR2 R: 5′-GAGGACTCGAGCTCAAGC-3′

### Additional oligos used in this study

U2AF2 shRNA1 F: 5′-ccggcgacgaggagtatgaggagatctcgagatctcctcatactcctcgtcgtttttg-3′

U2AF2 shRNA1 R: 5′-aattcaaaaacgacgaggagtatgaggagatctcgagatctcctcatactcctcgtc-3′

U2AF2 shRNA2 F: 5′-ccggtgcagattaaccaggacaagactcgagtcttgtcctggttaatctgcaattttg-3′

U2AF2 shRNA2 R: 5′-aattcaaaaatgcagattaaccaggacaagactcgagtcttgtcctggttaatctgc-3′

U2AF1 shRNA1 F: 5′-ccggccgacgtttagccagaccattctcgagaatggtctggctaaacgtcggttttt-3′

U2AF1 shRNA1 R: 5′-aattcaaaaaccgacgtttagccagaccattctcgagaatggtctgcctaaacgtcg-3′

Ctrl AMO: 5′-CCTCTTACCTCAGTTACAATTTATA-3′

U1 AMO: 5′-GGTATCTCCCCTGCCAGGTAAGTAT-3′

U2 AMO: 5′-TGATAAGAACAGATACTACACTTGA-3′

Ctrl Vivo-AMO: 5′-CCTCTTACCTCAGTTACAATTTATA-3′

*TP53* 5′ ss Vivo-AMO: 5′-TTCAGTCAGGAGCTTACCCAATCCA-3′

*TP53* 3′ ss Vivo-AMO: 5′-GTCTGGCTGCTGCAAGAGGAAAAGT-3′

### Statistics

All statistics were performed using the GraphPad Prism 10.2.1 software or RStudio. Statistical tests used are listed in the figure legends. Where applicable, unlabeled indicates p-value > 0.05, * indicates p-value ≤ 0.05, ** indicates p-value ≤ 0.01, *** indicates p-value ≤ 0.001, and **** indicates p-value ≤ 0.0001. For all boxplots, the Tukey method was used for error bars. For boxplots with more than two groups, statistical analyses were calculated using Kruskal-Wallis test with Dunn’s correction for multiple comparisons.

---

**Figure S1.**
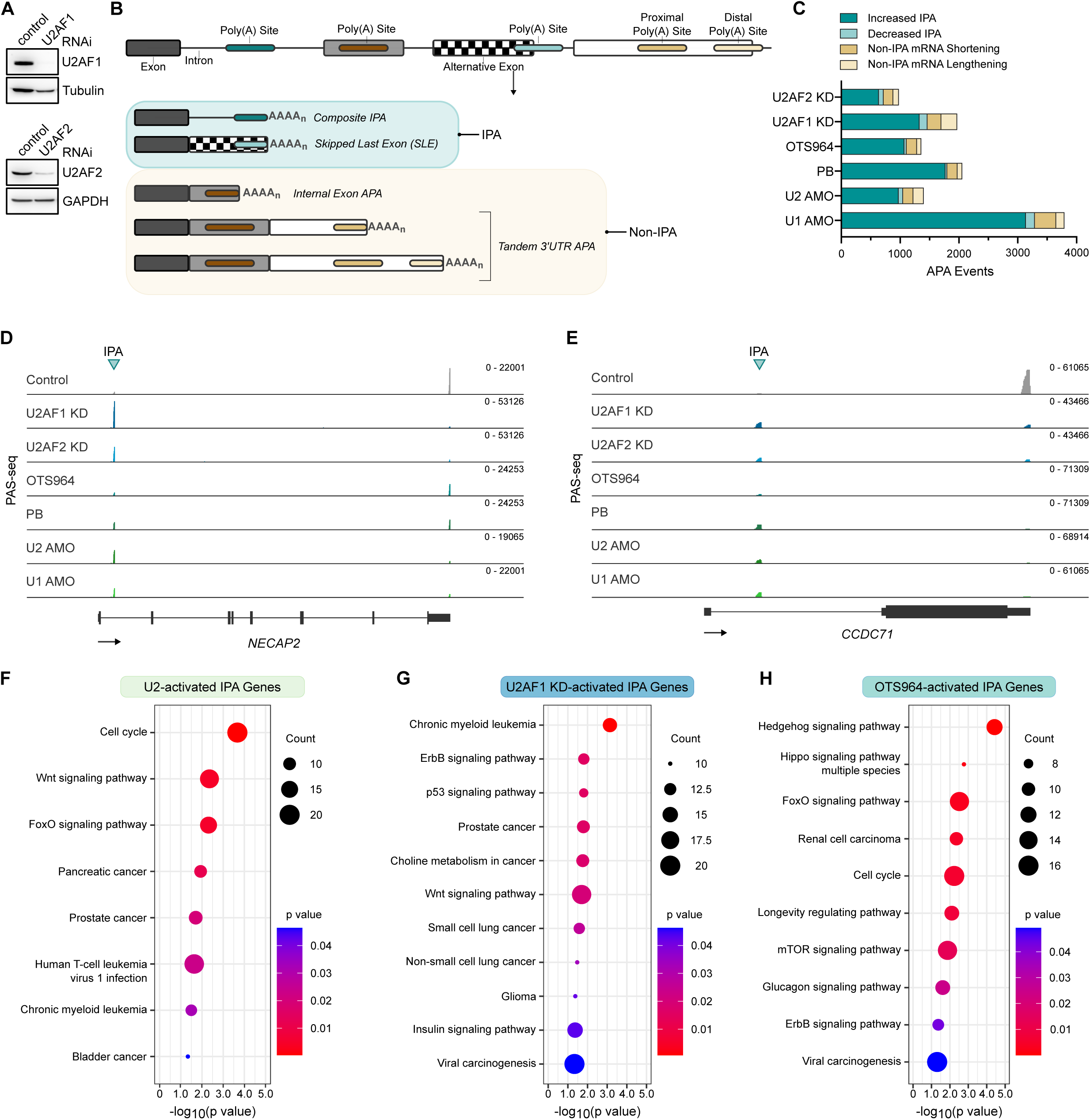
Global splicing inhibition activates shared and distinct intronic polyadenylation sites. **(A)** Western blot analysis of knockdown samples prior to PAS-seq to confirm depletion of U2AF1 or U2AF2. Tubulin and GAPDH were used as loading controls. **(B)** Schematic depicting the synthesis of IPA transcripts and non-IPA transcripts. IPA transcripts are produced when a pre-mRNA is cleaved and polyadenylated within the intron. For IPA transcripts that result from a skipped last exon (SLE), an alternative last exon is also produced by alternative splicing. For non-IPA transcripts, cleavage and polyadenylation may occur within an internal exon or within the terminal exon. **(C)** Stacked barplots depicting the frequency of genes with alternative polyadenylation site usage as classified by their dominant change following splicing inhibition treatments. **(D-E)** PAS-seq data tracks for the genes *NECAP2* (**D**) and *CCDC71* (**E**) following global splicing inhibition by depletion of U2AF1 or U2AF2 or treatment with OTS964, PB, U2 AMO, or U1 AMO in wild-type HEK293T cells. Each peak represents reads mapped to the 3′ end of mRNAs. **(F**-**H)** KEGG pathway analyses for activated IPA sites following treatment with U2 AMO (**F**), U2AF1 depletion (**G**), or OTS964 treatment (**H**). The terms related to cell development, growth and disease were collected. The terms in which p value < 0.05 and FDR < 0.05 were regarded as a significant functional annotation clustering.

**Figure S2.**
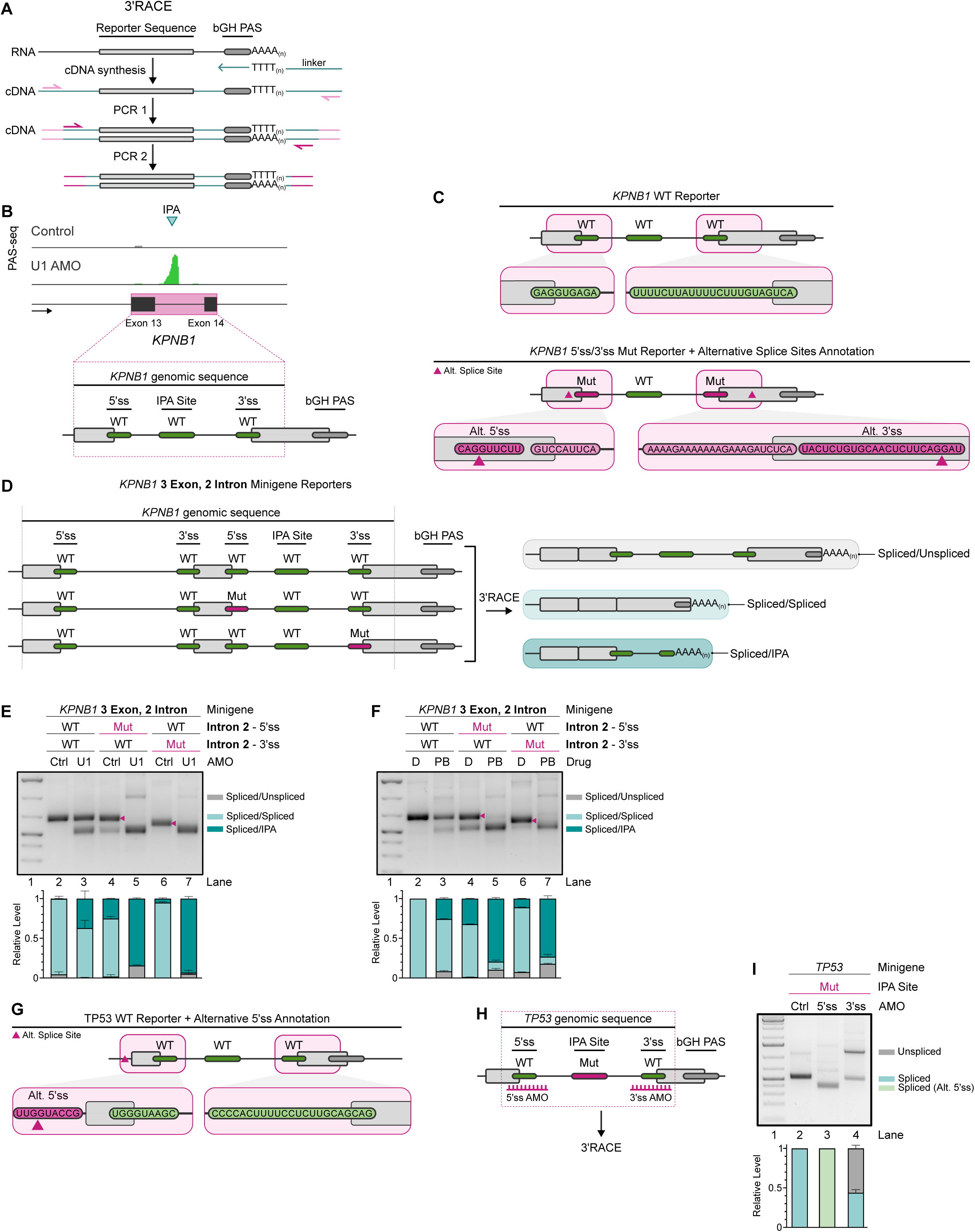
Additional characterization of intronic polyadenylation caused by gene-specific splicing inhibition. **(A)** Schematic of 3′RACE. **(B)** Top: PAS-seq data tracks for the gene *KPNB1* depicting an activated IPA site following global splicing inhibition by U1 AMO in wild-type HEK293T cells. The peak represents reads mapped to the 3′ end of mRNAs. Bottom: Schematic of the *KPNB1* genomic structure and sequences used to generate the *KPNB1* minigene reporter. **(C)** Top: Detailed schematic of the *KPNB1* WT minigene reporter with the WT 5′ and 3′ ss sequences included. Bottom: Detailed schematic of the *KPNB1* 5′ ss/3′ ss Mut reporter with the Mut 5′ ss and 3′ ss sequences included. The locations and sequences of a cryptic 5′ and 3′ ss, as detected by Sanger sequencing of 3′RACE products, are also included. **(D)** Schematic of *KPNB1* three-exon minigene reporter used for 3′ RACE. **(E - F)** 3′ RACE analysis of the *KPNB1* three-exon minigene reporter in control AMO and U1 AMO treated cells **(E)** or DMSO and PB-treated cells **(F)**. Top: Agarose gel used to resolve 3′RACE products. Bottom: Quantification of 3′RACE performed using a fragment analyzer (see Methods). Each stacked barplot includes the relative abundance of 3′ RACE products that correspond to spliced first intron and unspliced second intron (spliced/unspliced), both introns spliced (spliced/spliced), or a spliced first intron and IPA in the second intron (spliced/IPA). Data are presented as mean ± SD (*n* = 3). **(G)** Detailed schematic of the *TP53* minigene reporter with the WT 5′ and 3′ ss sequences included. The location and sequence of a cryptic 5′ ss, as detected by Sanger sequencing of 3′RACE products, located in the plasmid backbone is also included. **(H)** Schematic of the *TP53* minigene reporter in which the IPA site was mutated. **(I)** 3′ RACE analysis of the *TP53* IPA site Mut minigene reporter in control-treated, 5′ss AMO-treated, or 3′ss AMO-treated cells. Top: Agarose gel used to resolve 3′RACE products. Bottom: Quantification of 3′RACE performed using a fragment analyzer (see Methods). Each stacked barplot includes the relative abundance of unspliced, spliced, spliced using an alternative 5′ ss, or IPA 3′ RACE products in the specified sample. Data are presented as mean ± SD (*n* = 3).

**Figure S3.**
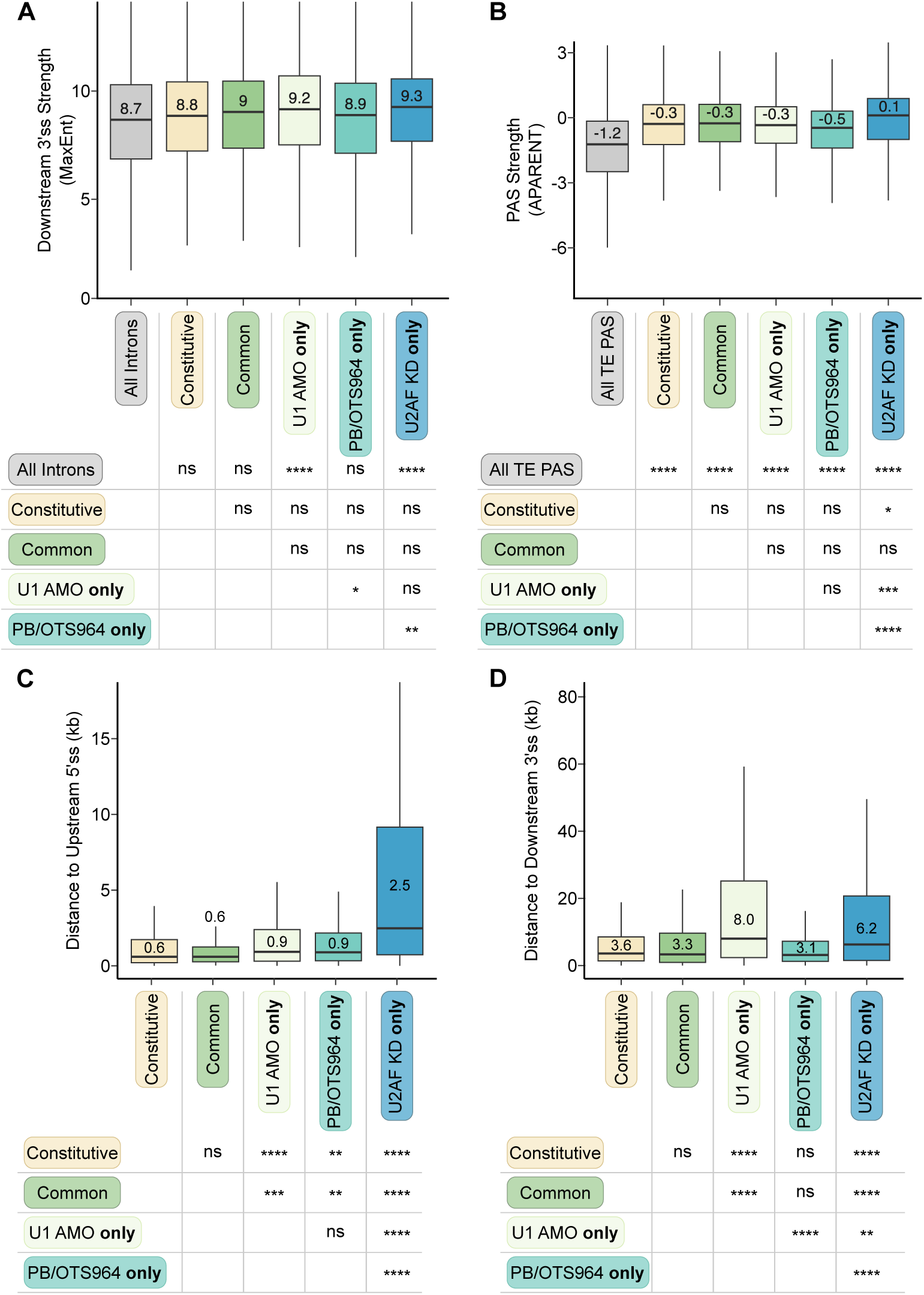
Sequence features of intronic polyadenylation sites activated by global splicing inhibition. **(A)** Boxplots depicting the strength of the nearest downstream 3′ ss as measured by MaxEnt for significantly activated IPA sites following each global splicing inhibition treatment. Statistical analysis was calculated by Kruskal-Wallis test. ∗p value ≤ 0.05, ∗∗p value ≤ 0.01, ∗∗∗p value ≤ 0.001, ∗∗∗∗: p-value ≤ 0.0001. **(B)** Boxplots depicting the PAS strength as measured by APARENT of significantly activated IPA sites following each global splicing inhibition treatment. Statistical analysis was calculated by Kruskal-Wallis test. ∗p value ≤ 0.05, ∗∗p value ≤ 0.01, ∗∗∗p value ≤ 0.001, ∗∗∗∗: p-value ≤ 0.0001. **(C)** Boxplots depicting the distance between the nearest upstream 5′ ss and the activated IPA site for significantly activated IPA sites following each global splicing inhibition treatment. Statistical analysis was calculated by Kruskal-Wallis test. ∗p value ≤ 0.05, ∗∗p value ≤ 0.01, ∗∗∗p value ≤ 0.001, ∗∗∗∗: p-value ≤ 0.0001. **(D)** Boxplots depicting the distance between the nearest downstream 3′ ss and the activated IPA site for significantly activated IPA sites following each global splicing inhibition treatment. Statistical analysis was calculated by Kruskal-Wallis test. ∗p value ≤ 0.05, ∗∗p value ≤ 0.01, ∗∗∗p value ≤ 0.001, ∗∗∗∗: p-value ≤ 0.0001.

**Figure S4.**
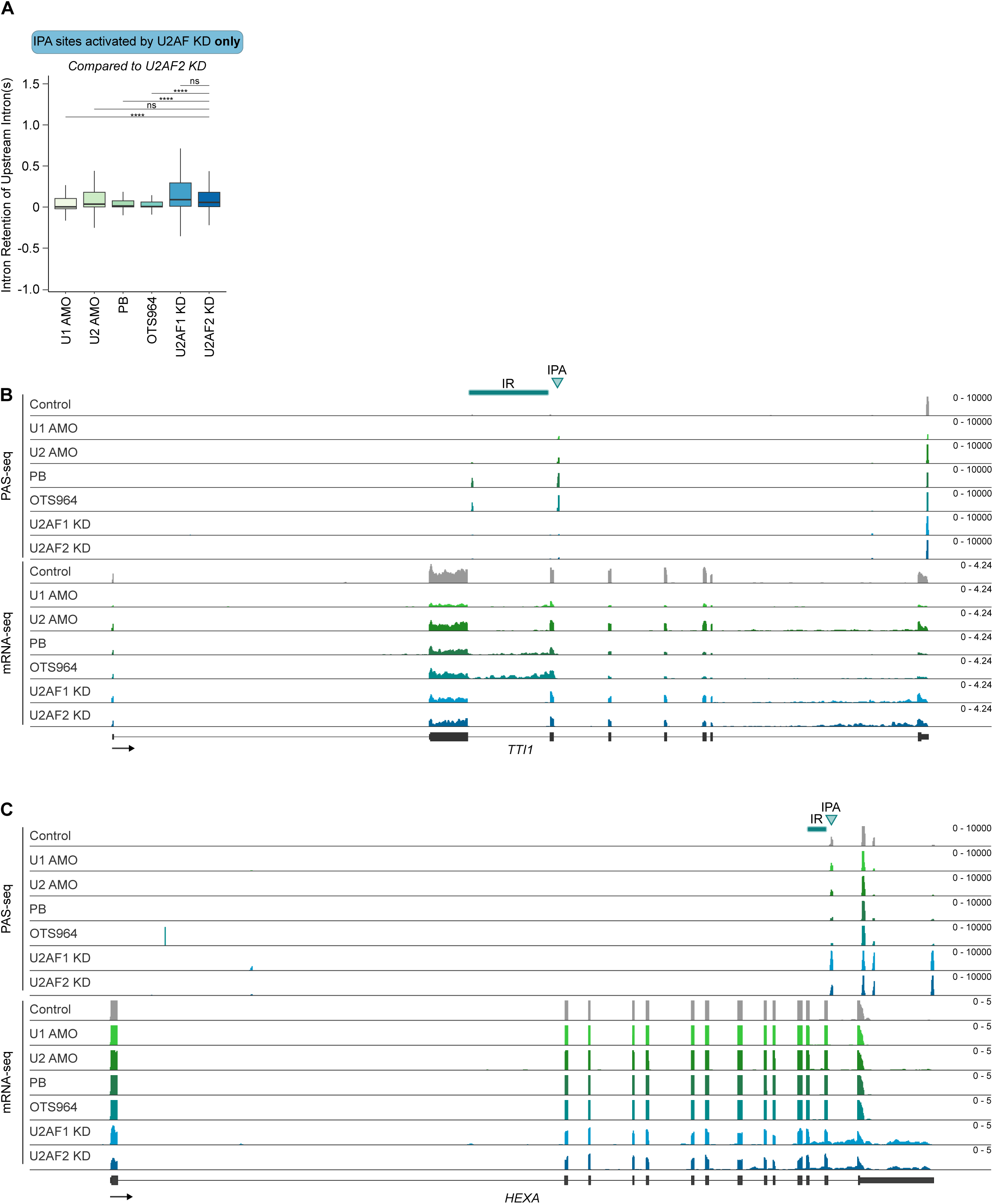
Additional characterization of how splicing inhibition affects splicing and IPA in a coordinated manner. **(A)** Boxplots depicting the IR ratio of introns upstream of IPA sites that are exclusively activated by U2AF KD. Statistical analysis was calculated by Kruskal-Wallis test and comparisons are present between U2AF2 KD and all other groups. ∗p value ≤ 0.05, ∗∗p value ≤ 0.01, ∗∗∗p value ≤ 0.001, ∗∗∗∗: p-value ≤ 0.0001. **(B - C)** PAS-seq and mRNA-seq data tracks for the gene *TTI1* **(B)** and *HEXA* **(C)** following the indicated splicing inhibition treatments. Each peak in PAS-seq represents reads mapped to the 3′ end of mRNAs. IR is depicted by a bar and IPA is indicated by an arrow. The y-axis of mRNA-seq and PAS-seq for the gene *HEXA* **(C)** is truncated to better display intronic signal.

**Figure S5.**
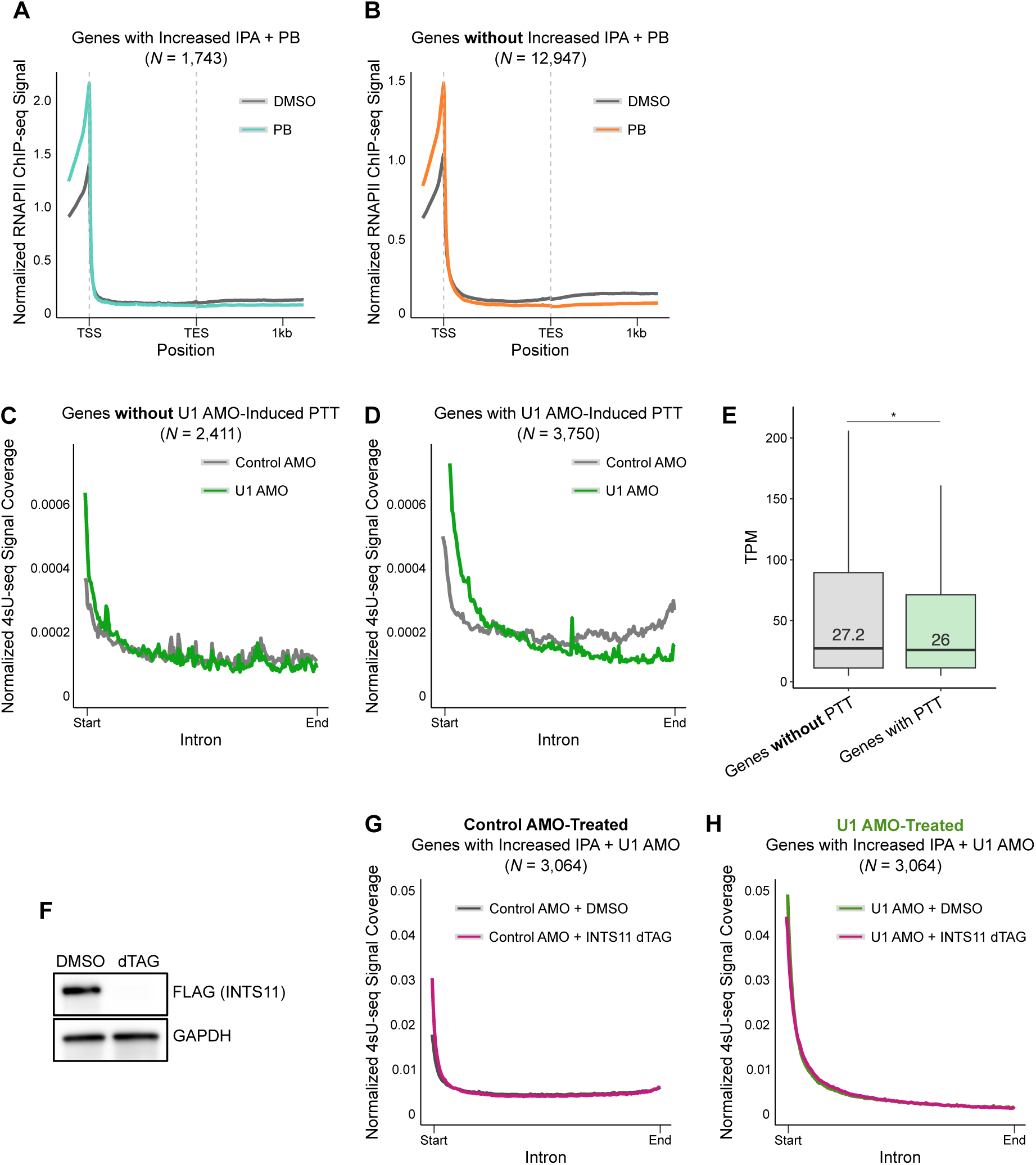
INTS11 is not required for U1 AMO-induced premature transcription termination. **(A)** Metagene plot depicting the RNAPII ChIP-seq signal in DMSO-treated cells (gray line) or PB-treated cells (blue line) for all expressed genes that contained an activated IPA site following PB treatment as detected by PAS-seq (*N* = 1,743). The x-axis depicts a composite intron in which all introns of each gene were combined to determine the relative position of ChIP-seq signal. **(B)** Metagene plot depicting the RNAPII ChIP-seq signal in DMSO-treated cells (gray line) or PB-treated cells (orange line) for all expressed genes that did not contain an activated IPA site following PB treatment as detected by PAS-seq (*N* = 12,947). The x-axis depicts a composite intron in which all introns of each gene were combined to determine the relative position of ChIP-seq signal. **(C)** Metagene plot depicting the 4sU-seq signal in HEK293T cells that were treated with control AMO (gray line) or U1 AMO (green line) for all expressed genes that did not exhibit U1 AMO-induced PTT (*N* = 2,411). The x-axis depicts a composite intron in which all introns of each gene were combined to determine the relative position of 4sU-seq signal. **(D)** Metagene plot depicting the 4sU-seq signal in HEK293T cells that were treated with control AMO (gray line) or U1 AMO (green line) for all expressed genes that exhibited U1 AMO-induced PTT (*N* = 3,750). The x-axis depicts a composite intron in which all introns of each gene were combined to determine the relative position of 4sU-seq signal. **(E)** Boxplot depicting the transcripts per million (TPM) for genes without U1 AMO-induced PTT (*N* = 2,411) and genes with U1 AMO-induced PTT (*N* = 3,750). Statistical analysis was calculated by t-test. ∗p value ≤ 0.05 **(F)** Western blot analysis of INTS11 and GAPDH (loading control) levels in the INTS11 degron cell line that were DMSO-treated or dTAG-treated for 16 hours prior to 4sU-seq. **(G)** Metagene plot depicting the 4sU-seq signal in INTS11 degron cells that were treated with control AMO and DMSO (gray line) or control AMO and dTAG (pink line) for all expressed genes that contained an activated IPA site following U1 AMO treatment as detected by PAS-seq (*N* = 3,064). The x-axis depicts a composite intron in which all introns of each gene were combined to determine the relative position of 4sU-seq signal. **(H)** Metagene plot depicting the 4sU-seq signal in INTS11 degron cells that were treated with U1 AMO and DMSO (green line) or U1 AMO and dTAG (pink line) for all expressed genes that contained an activated IPA site following U1 AMO treatment as detected by PAS-seq (*N* = 3,064). The x-axis depicts a composite intron in which all introns of each gene were combined to determine the relative position of 4sU-seq signal.

**Figure S6.**
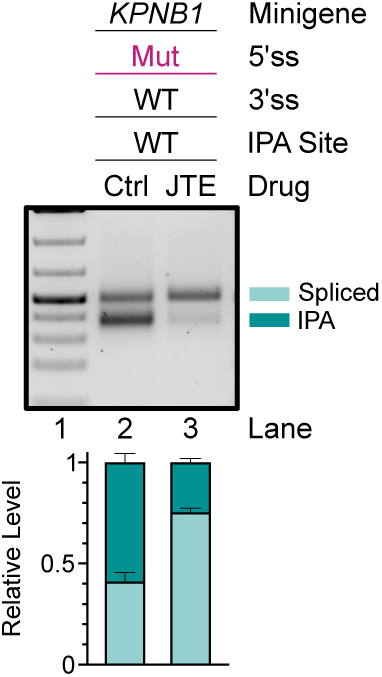
Inhibition of pre-mRNA 3′ processing increases splicing efficiency in a reporter assay. 3′ RACE analysis of the *KPNB1* 5′ ss Mut minigene reporter in control- or JTE-607-treated cells. Top: Agarose gel used to resolve 3′RACE products. Bottom: Quantification of 3′RACE performed using a fragment analyzer (see Methods). Each stacked barplot includes the relative abundance of unspliced, spliced, spliced using an alternative 5′ ss, or IPA 3′ RACE products in the specified sample. Data are presented as mean ± SD (*n* = 3).

## Notes

### Competing Interest Statement

The authors have declared no competing interest.

